# Kinetic modeling of leucine-mediated signaling and protein metabolism in human skeletal muscle

**DOI:** 10.1101/2023.06.10.544441

**Authors:** Taylor J. McColl, David C. Clarke

## Abstract

Skeletal muscle protein levels are governed by the relative rates of muscle protein synthesis (MPS) and breakdown (MPB). The mechanisms controlling these rates are complex and their integrated behaviors are challenging to study through experiments alone. The purpose of this study was to develop and analyze a kinetic model of leucine-mediated mTOR signaling and protein metabolism in the skeletal muscle of young adults. Our model amalgamates published cellular-level models of the IRS1-PI3K-Akt-mTORC1 signaling system and of skeletal-muscle leucine kinetics with physiological-level models of leucine digestion and transport and insulin dynamics. The model satisfactorily predicts experimental data from diverse leucine feeding protocols. Model analysis revealed that basal levels of p70S6K are a primary determinant of MPS, insulin signaling substantially affects muscle net protein balance via its effects on MPB, and p70S6K-mediated feedback of mTORC1 signaling reduces MPS in a dose-dependent manner.

## Introduction

Skeletal muscle enables locomotion, metabolic regulation^1^, and physical performance^2, 3^. Skeletal muscle function depends largely on its mass^4^, such that maintaining or increasing skeletal muscle mass benefits human health and quality of life. Conversely, the loss of skeletal muscle mass in disease states such as sarcopenia negatively impacts health. The prevalence of sarcopenia in older adults (≥60 years) ranges from 9% to 51% depending on factors such as gender, age, pathological condition, and diagnostic criteria^5–7^. Those that are sarcopenic are hospitalized more frequently^8^ and engender costlier care^9–11^. Therefore, deeper understanding of skeletal muscle mass regulation and effective therapies that promote skeletal muscle mass are sought.

Skeletal muscle mass is determined predominantly by muscle protein levels. Muscle protein levels are governed by the relative rates of muscle protein synthesis (MPS) and muscle protein breakdown (MPB). MPS rates are highly sensitive to amino acid concentrations, in particular leucine^12–15^, which promotes protein translation initiation and elongation^16–18^. Translation initiation is considered the “rate-limiting step” for protein biosynthesis^19^. However, feeding only promotes MPS for a finite duration, even with sustained amino acid availability in the blood plasma^20, 21^. MPB rates vary less than those of MPS^21, 22^. Thus, the daily fluctuations of MPS above and below MPB, coinciding with fed and fasted states, respectively, are thought to determine overall muscle protein balance. Sustaining a neutral or positive protein balance requires that amino acids be ingested at regular intervals throughout the day^20, 21^.

Muscle protein balance is controlled by three principal mechanisms: 1) hormones (e.g., insulin and insulin-like growth factors, 2) nutrition (e.g., amino acids), and 3) mechanical load (e.g., resistance training)^18^. These factors converge on the protein kinase mechanistic target of rapamycin complex 1 (mTORC1), which integrates the stimuli to control MPS. Of these factors, the amino acid leucine activates mTORC1 through both leucine- and insulin-dependent signaling pathways, because leucine also promotes insulin secretion^23, 24^. Leucine is sensed by Sestrin2, which activates the Ragulator-Rag complex and ultimately activates the mTORC1 kinase^25–27^. Insulin activates mTORC1 through a signaling cascade that involves insulin receptor (IR), insulin receptor substrate 1 (IRS1), phosphoinositide 3-kinase (PI3K), 3-phosphoinositide dependent pprotein kinase-1 (PDK1), protein kinase B (Akt), and the tuberous sclerosis protein complex 1/2 (TSC1/2)^28–30^. Activated mTORC1 phosphorylates p70 ribosomal protein S6 kinase (p70S6K) and eukaryotic initiation factor 4E-binding protein 1 (4EBP1), which promote translation initiation^19, 31^ and subsequently MPS. Insulin signaling reduces MPB via anticatabolic effects^32–35^, but the mechanisms remain incompletely understood. It is currently thought that insulin modulates MPB through the ubiquitin-proteasomal system and lysosomal autophagy pathway, which act via interactions between Akt/forkhead box O3 (FoxO3) and mTORC1/unc-51 like autophagy activating kinase 1 (ULK1), respectively^22, 36, 37^.

Diverse experimental methods are used to study the physiological and cellular mechanisms of feeding-induced skeletal muscle protein metabolism. Measurements of blood plasma are used to infer the secretion rates of hormones (e.g., insulin) and the rates of absorption of relevant amino acids. The specific amino acids measured vary depending on the ingested solution, but individual amino acids (e.g., leucine, phenylalanine) and general classes of amino acids (e.g., essential amino acids (EAA), branched chained amino acids) are commonly assessed. Measurements of biopsied muscle are used to assess the post-translational modifications of key proteins in the mTOR signaling cascade (e.g., phospho-Akt, phospho-p70S6K). These phosphorylated proteins are used to infer anabolic signaling, e.g., increased phospho-p70S6K levels imply accelerated MPS. Stable isotope tracers are the primary method used to assess MPS rates. Stable isotopes are added to the nutritional solution consumed by the study participants to determine the rates of incorporation of traced amino acids into skeletal muscle. The enrichment of isotopes in the muscle tissue, measured via serial muscle biopsy samples, is used to estimate the muscle protein synthesized during a specified duration [i.e., the fractional synthetic rate (FSR)]. Data regarding stable isotope tracer enrichment in arterial plasma, venous plasma, and muscle are used to estimate the parameters of models of amino acid dynamics (e.g., the 3-pool model^38, 39^). These models enable the inference of transmembrane amino acid transport and muscle protein kinetics across the whole-muscle^38, 39^. While the mechanisms controlling muscle protein metabolism have been identified and characterized, how these mechanisms act as part of an integrated, multi-scale system remain poorly understood. Accordingly, the relative contributions of protein digestion and transport, hormonal signaling, intracellular anabolic signaling, and amino acid metabolism to overall muscle protein metabolism have yet to be quantified.

Mathematical models are powerful tools for analyzing complex multiscale biological systems. Models enable the integration of chemical and physical principles, prior knowledge, and experimental data in a coherent framework^40^. Mathematical models have been developed for aspects of protein metabolism and anabolic signaling. For example, the mechanisms controlling the ultradian secretion of insulin from the pancreas have been studied using models that incorporate the negative feedback loops between insulin and glucose^41, 42^. Models of amino acid dynamics have expanded upon the previously mentioned 3-pool model and include a six-compartment model of intracellular muscle kinetics of leucine and its transamination product ⍺-ketoisocaproic acid (KIC) across the human forearm^43^ and a ten-compartment model of human protein kinetics incorporating compartments for leucine, KIC, and bicarbonate^44^. Models of mTORC1 signaling have investigated the control of mTOR by amino acids and insulin^45^, the control of mTOR complex 2 (mTORC2) in response to amino acid and insulin stimulation^46^, the influence of AMP-activated protein kinase in response to amino acid and insulin stimulation on downstream mTORC1 activity^47^, and novel amino acid inputs to the mTOR network^48^. However, an integrated model of muscle protein metabolism combining insulin secretion, amino acid dynamics, and mTORC1 signaling has yet to be proposed.

The purpose of this study was to develop and analyze a kinetic model of skeletal muscle protein metabolism. The model focused on leucine dynamics because it is the primary amino acid governing muscle protein synthesis^12, 49–51^. A compartmental kinetic model is proposed that simulates leucine incorporation into skeletal muscle following ingestion of leucine downstream of core signaling and metabolic processes. The model satisfactorily simulates leucine-mediated signaling and protein metabolism dynamics following bolus and pulsatile feeding. We analyzed the model via simulations to demonstrate that the basal levels of p70S6K are a primary determinant of MPS, more so than changes in phospho-p70S6K levels, insulin signaling substantially affects net muscle protein balance through inhibition of MPB, and p70S6K-mediated feedback of mTORC1 signaling reduces MPS in a dose-dependent manner.

## Methods

### Model construction – system definition and simplifying assumptions

We defined our system to consist of four compartments: 1) stomach, 2) gut, 3) blood plasma/interstitial fluid, and 4) skeletal muscle. The system is stimulated through leucine feeding, in which fed leucine travels through a digestive system input module and is absorbed into the blood plasma compartment. The blood plasma compartment subsequently drives the cellular signaling and leucine dynamics in the skeletal muscle compartment. Each compartment is assumed to behave as a well-stirred tank reactor, and the modelled proteins were assumed to exist in sufficient concentrations to behave deterministically. These simplifying assumptions enabled us to use ordinary differential equations (ODEs) as the model’s mathematical framework.

### Model construction – topology and biochemistry considerations

Our model features four modules: 1) mTOR signaling module, 2) leucine kinetic module, 3) digestive system module, and 4) insulin dynamics module.

#### The mTOR signaling module

The starting framework for the mTOR module was the model of Dalle Pezze et al.^46^. The Dalle Pezze et al.^46^ model simulates the insulin signaling dynamics propagating across the IR/Akt/mTORC1/p70S6K axis. The model also included an independent pathway for amino-acid-stimulated mTORC1 activity. We replicated the model representing their “hypothesis four” by using their ODEs, parameter values, and initial conditions. We were able to replicate their model outputs after slightly adjusting two of the parameter values. This result supports the correctness of the code upon which subsequent model iterations were developed.

The Dalle Pezze et al.^46^ model featured three shortcomings that we addressed to fulfill our modelling objectives. First, the model inputs (insulin and amino acids) were set as constant values for the simulation duration^46^, which is inconsistent with the fact that both are dynamic, especially following feeding. Second, the model was developed using data from HeLa cells, such that we needed to adapt the model to human skeletal muscle. Third, the IRS1-PI3K module contained a degradation (“sink”) term. This term simplifies the IRS1-PI3K module, so we replaced the sink term with a more explicit representation of the signaling mechanisms. Specifically, we incorporated three IRS1 species (IRS1, p-IRS1^Y^, and p-IRS1^S^) and specified an association reaction between p-IRS1^Y^ and PI3K to form the p-IRS1^Y^-PI3K complex that promotes PDK1 phosphorylation. Several serine residues on IRS1 can be phosphorylated, which we designated p-IRS^S^ to collectively represent the species, because each inhibits downstream pathways in a similar manner.

#### The leucine kinetic module

We developed functions to simulate the dynamic amino acid and insulin inputs to the model. Leucine is the amino acid that most potently stimulates mTORC1 activity^16–18^, such that we aimed to locate models of leucine kinetics that could replace the amino acid input from the Dalle Pezze et al.^46^ model. We located the Tessari et al.^43^ model of leucine kinetics, which used six compartments to simulate the dynamics of leucine across the human forearm. We developed a working model of leucine dynamics using the rate equations and flow rates provided in Tessari et al.^43^, but the model needed to be modified in several ways to facilitate its integration with the mTOR signaling module. First, we assumed that the blood plasma/interstitial fluid compartment in our model was a well-mixed reactor (i.e., a mixture of arterial and venous blood) so we removed the venous leucine and KIC compartments. Removal of these compartments reduced the model from six to four compartments. The differences in concentrations between the arterial and venous compartments of leucine and KIC were 0.1 and 0.7 μmol/L, respectively, which we considered negligible (i.e., arterial leucine = 122.9 μmol/L, venous leucine = 122.8 μmol/L; arterial KIC = 25.8 μmol/L, venous KIC = 26.5 μmol/L).

Second, Tessari et al.^43^ reported transport rates rather than rate constants, such that we calculated the latter using the following equation:

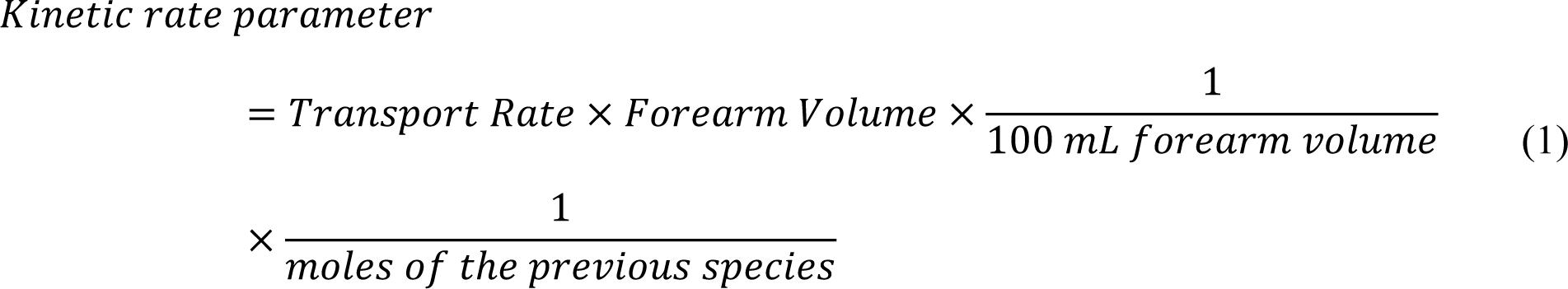

where the transport rate is in units of *nmol* · *min*^−1^ · 100 *mL*^−1^, forearm volume is in units of *mL*, and moles of the previous species is in *nmol*.

#### The digestive system module

We added a digestive system module that operates as an input function to simulate the dynamics of leucine ingestion and absorption in the blood. This module included two compartments: stomach and gut, and two degradation terms: excretion of leucine via the gut (i.e., feces) and first-pass splanchnic extraction. The parameters for this module reflect the gastric emptying and absorption rate of leucine when ingested in a low-volume, low-caloric solution in the fasted state^52^ and were informed using the true ileal digestibility of leucine (∼90%)^53^ and the first-pass splanchnic extraction of amino acids in young adults (23-29%)^54, 55^.

#### The insulin secretion module

We used the previously validated model of Sturis et al.^41^ to simulate physiologically realistic insulin dynamics. Insulin is secreted from the pancreas in an oscillatory pattern with a period of approximately 120 minutes (i.e., ultradian)^41^.The Sturis et al.^41^ model mimics the ultradian oscillations of insulin secretion by featuring four negative feedback loops: 1) elevated glucose concentrations stimulate insulin secretion which reduces glucose production, 2) elevated glucose concentrations stimulate insulin secretion which promotes glucose utilization, 3) glucose inhibits further glucose production, and 4) glucose promotes further glucose utilization. The Sturis et al.^41^ model includes three compartments (plasma insulin, intercellular insulin, plasma glucose) and three variables that represent the delay between plasma insulin and its effect on hepatic glucose production.

We modified the Sturis et al.^41^ model in two ways to integrate it with our model. First, all glucose- or insulin-specific parameters (i.e., any parameters with units of mg or mU, respectively) were converted to units of moles using the molecular mass of glucose (180.156 g/mol) or the insulin unit conversion factor provided by Sturis et al.^41^ (1 mU insulin ≅ 6.67 pmol). Second, we set the basal glucose infusion rate in the Sturis et al.^41^ model to 75 mg/min to maintain basal plasma insulin concentrations at approximately 26 pmol/L (i.e., the concentration of plasma insulin from the calibration data set).

#### Integration of modules

Having successfully replicated the mTOR signaling and insulin secretion modules and having formed working modules for leucine kinetics and the digestive system, we integrated the four modules through four links. First, intracellular leucine replaced the constant amino acid stimulus from the Dalle Pezze et al.^46^ model as the activating stimulus for mTORC1. Second, we added p70S6K^T389^ as a controller of intracellular leucine incorporation into skeletal muscle protein (i.e., protein synthesis) because phospho-p70S6K controls protein synthesis^31, 56^. Third, leucine is an insulin secretagogue^23^, such that we added a link to simulate leucine-mediated insulin secretion. Lastly, insulin regulates skeletal muscle mass primarily by reducing MPB^32^. We were unable to find from the literature the precise mechanism by which insulin signaling controls MPB, such that we chose p-IR_ß_^Y^ as the effector of insulin signaling within the skeletal muscle cell to inhibit MPB (i.e., elevated p-IR_ß_^Y^ reduces MPB, whereas reduced p-IR_ß_^Y^ increases MPB).

After integrating the modules, we expanded the model topology to represent the current state of knowledge in the literature while fostering model parsimony. We made four modifications to the model:

1. Akt is an AGC kinase that must be phosphorylated twice for full activity^46^. PDK1 and a PDK2 phosphorylate the Thr308 and Ser473 residues of Akt, respectively. mTORC2 is a *bona fide* PDK2 that phosphorylates the Ser473 residue of Akt^46^. However, Dalle Pezze et al.^46^ were unable to reproduce experimental data for p-Akt^S473^ with mTORC2 alone, such that they introduced an additional PDK2 component to resolve the model output. However, the literature suggests that mTORC2 is the primary PDK2 that drives p-Akt^S473^ ^57, 58^, such that we felt justified in removing the additional PDK2 species.
2. We expanded the Akt module to include all possible combinations of its phosphorylation states (i.e., Akt^T308^, Akt^S473^, Akt^S473,T308^)^28^.
3. Phospho-p70S6K inhibits mTORC1 activity through a negative feedback loop that results in the phosphorylation of mTORC1 at Ser2448, a residue that resides in the mTOR negative regulatory domain^59–62^. However, the Dalle Pezze et al.^46^ model used phospho-mTORC1^S2448^ as a marker of mTORC1 activity. We therefore redefined the mTORC1 species as either *active* or *inactive* and included a negative feedback loop from phospho-p70S6K to mTORC1.
4. For the sake of model parsimony, we removed the proline-rich Akt substrate of 40 kDa (PRAS40) because of a lack of experimental data for model fitting and because PRAS40 did not influence other model components.

A schematic diagram of the final model topology is presented in Figure 1. The model includes four defined compartments (stomach, gut, blood plasma, and skeletal muscle cells), 34 species (11 proteins, 13 post-translationally modified proteins, and 10 pools), 63 kinetic parameters (60 adjustable, three constrained), and three delay parameters between insulin and glucose production as per the Sturis et al.^41^ model (Table S1). All equations in our model were assumed to follow mass-action kinetics. The overall model consisted of 37 nonlinear ODEs that describe the rate of change of the number of moles of each molecular species within the indicated compartment (Tables S2-3). The number of moles of each species are determined by the sum of the reactions that generate and consume each species, which was expressed mathematically according to Equation 2.

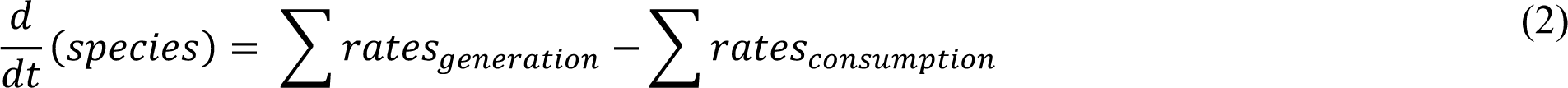

**Figure 1.**
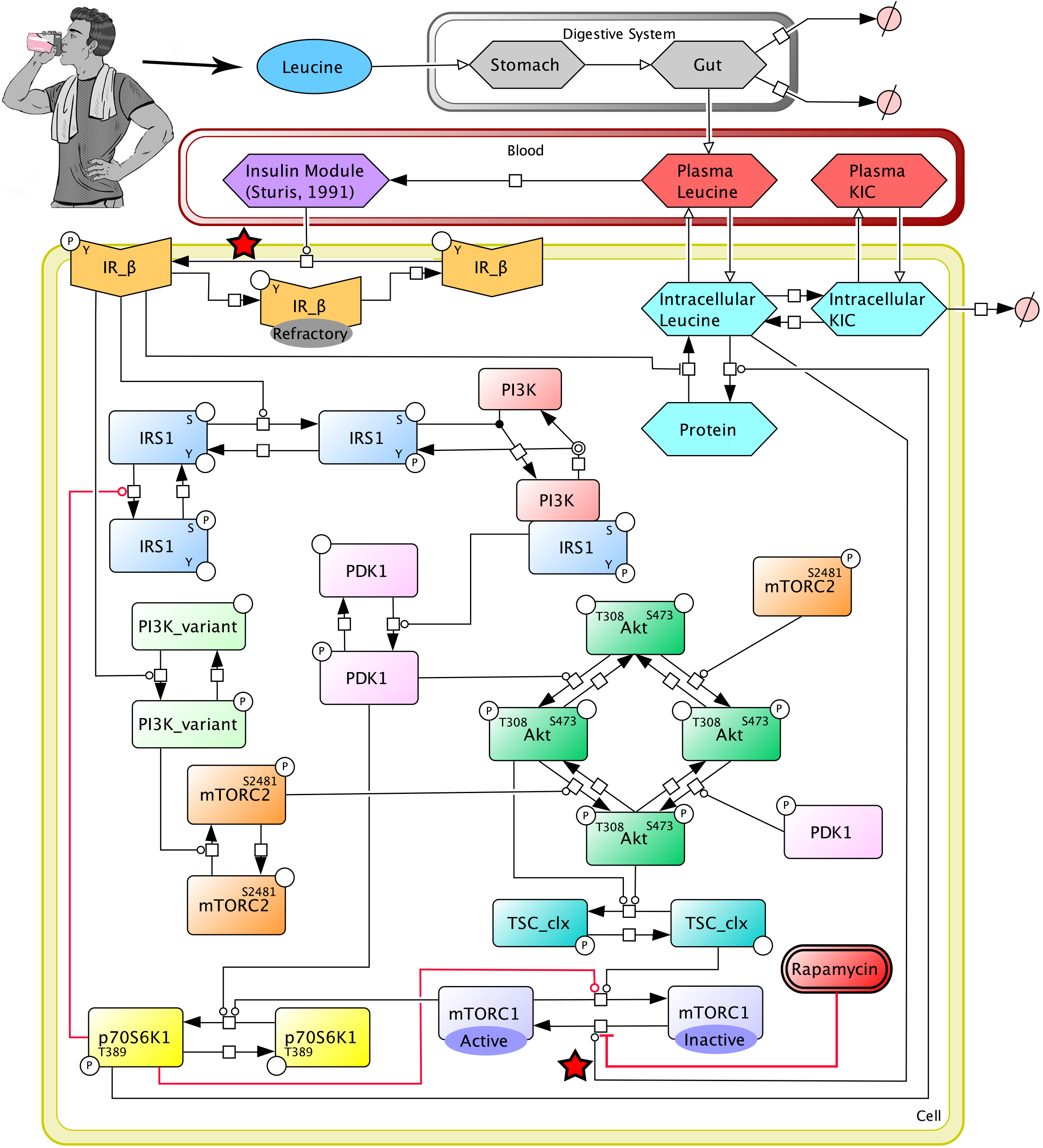
Reaction diagram of the model. Reaction diagram for the kinetic model of leucine-mediated signaling and protein metabolism. mTORC2^S2481^ and PDK1^P^ are represented as independent species for the phosphorylation of Akt^S473^ and Akt^S473,T308^, respectively, but both are subject to the same control as the integrated species. The insulin module is presented as a single species in the model diagram, but it is expanded in the model code to include the three species and four negative feedback loops outlined by Sturis et al.^41^. The transfer of masses is denoted by open headed arrows. Chemical reactions are denoted by solid headed arrows. Inhibition is denoted by a flat-headed line. Red stars denote the locations of simulated signaling knockdown. P = phospho residue, S = serine, T = threonine, Y = tyrosine.

All model runs were initiated with a burn-in period of 300 minutes to allow all model species to reach steady state. A plasma leucine infusion rate of 2.5 mg/min was used during the equilibrium period to maintain the physiological concentrations of species in the leucine module.

### Model calibration

Our model was calibrated similar to the procedure outlined in Zhao et al.^63^. First, we changed the initial conditions of the proteins to values representative of human skeletal muscle (Table S4). The concentration for plasma insulin was set to 28 pM^64^. The initial concentration for plasma leucine was set to 121 μM^65^. The initial value for plasma α-ketoisocaproate (KIC) was set to 30.6 μM^43^. The initial concentration for intracellular leucine was set to 123 μM based on the observation that intracellular leucine is 25% higher than plasma leucine^44^. The initial concentration of total leucine bound to skeletal muscle protein was set to 134 mM using the approach in Wolfe et al.^53^. Skeletal muscle mass was assumed to constitute 40% body weight and approximately 20% of that mass is assumed to be muscle protein. Assuming an average body mass of 72.1 kg, then the assumed skeletal muscle protein mass would be ∼5.8 kg (72.1 kg × 40% skeletal muscle × 20% muscle protein). We assumed that leucine comprises 8% of muscle protein mass^44^, such that there is ∼460 g of leucine content in skeletal muscle. We were unable to find experimentally measured concentrations for intracellular KIC, so we set its initial value to 0 M, allowed the species to reach a steady-state value during the equilibration phase of the model simulations, and used the value that was reached at the end of the equilibration phase for all subsequent model simulations (11.5 μM).

We searched the literature to obtain plausible ranges for the initial concentrations of the insulin signaling proteins, but we were unable to locate their concentrations in human skeletal muscle cells. However, we found copy number values for the insulin signaling proteins from quantitative proteomic studies using HeLa^66^ and U2OS^67^ cell lines (Table S4). The cell volumes used for the calculation of concentration in HeLa and U2OS cells were 2.6 × 10^-12^ L and 4.0 × 10^-12^ L, respectively^67, 68^. Given the consistency of the concentrations in HeLa and U2OS cells, and given that the cell volume of a human skeletal muscle cell line (TE671RD) is similar to the HeLa and U2OS cell lines (2.8 × 10^-12^ L)^69^, we assumed that the concentrations obtained in HeLa and U2OS cells are representative of those in human skeletal muscle. The initial concentrations of all phosphorylated proteins in the Dalle Pezze^46^ model were set to zero; however, a proportion of proteins exist in the phosphorylated state even at rest, such that we set the initial concentrations of phospho-proteins to 20% of the non-phosphorylated values.

Once the initial conditions of all proteins were calibrated to values representative of those in human skeletal muscle, we converted all concentrations to moles to allow us to simulate the rate of change of particles for each molecular species (i.e., we factored out the volume of the respective compartment). Converting species from concentrations to moles allowed us to ignore the influence of different compartment volumes within the model (i.e., movement of molecules across compartments of different volumes have disproportionate concentration changes). To convert species from concentrations to moles, we needed to estimate the volumes of each compartment. We used general subject characteristics from Caucasian males to calculate the compartment volumes muscle (age: 41.9 years old, mass: 72.1 kg, height: 1.73 m, resistance index (height^2^/Ω): 57.9 cm^2^/Ω)^70^. We calculated blood plasma volume using the Nadler formula for men^71^ (Equation 6):

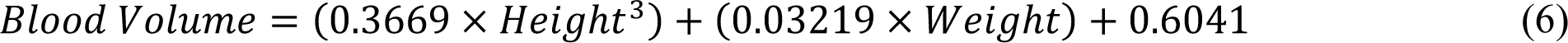

where blood volume is given in liters and height and weight are in units of meters and kilograms, respectively. We estimated the skeletal muscle volume by first estimating total skeletal muscle mass using the regression equation developed by Janssen et al.^70^ (Equation 7):

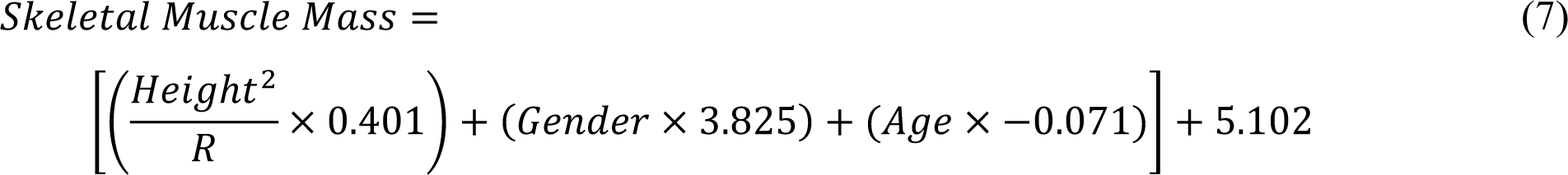

where R is the bioelectrical impedance analysis resistance in Ohms, height is in centimeters, age is in years, and gender is given a value of 1 or 0 for men and women, respectively. We then used the skeletal muscle mass to calculate the volume of skeletal muscle using the density value for mammalian skeletal muscle (1.112 g/mL)^72^.

Next, we adjusted the kinetic rate parameters (Table S5). To calibrate the kinetic rate parameters, we curated experimental time course data of eight readouts following insulin or leucine stimulation: plasma insulin^65^, plasma leucine^65^, intracellular leucine^73^, phospho-IR^74^, phospho-Akt^S473^, phospho-p70S6K1^T389^, and parameters from the three-pool model of leg amino acid kinetics [i.e., F_m,a_: inward amino acid transport from artery to muscle; F_m,0_: intracellular amino acid appearance from endogenous sources (i.e., proteolysis, de novo synthesis)]^65^. The phospho-Akt^S473^ and phospho-p70S6K1^T389^ time-courses were obtained by meta-analyzing studies that measured the respective protein in non-exercised, young adults following leucine ingestion (further details provided in the Comprehensive Model Development Procedure). Where necessary, we supplemented the muscle-specific data in response to leucine feeding with time courses following whey protein feeding (plasma KIC^75^) and from other cell types, including L6 myotubes (Akt^T308^ ^76^), and 3T3-L1 adipocyte-like cells [insulin receptor substrate 1 (IRS1)^77^]. We used these data to manually tune the parameter values to achieve a reasonable visual fit. We then used numerical optimization (“fmincon” and “GlobalSearch” functions in MATLAB) to fit the model parameters to quantitative time-courses of plasma insulin, plasma leucine, F_m,a_, and F_m,0_, to semi-quantitative time-courses (i.e., immunoblot data) of phospho-Akt^S473^ and phospho-p70S6K^T389^, and FSR data in response to a 3.5-gram bolus of leucine (Table S6). We used FSR data^65^ in the model calibration as a measure of the total MPS response (i.e., total grams of leucine synthesized into skeletal muscle over the intervention duration). We simulated the total MPS response in the model by measuring the area under the curve (AUC; “cumtrapz” function in MATLAB) of the MPS reaction (r15), we then converted the value from total moles over the intervention period to total grams using the leucine molar mass. We used the method in Wolfe et al.^53^ to convert experimentally measured FSR to total grams of leucine synthesized over the intervention period (Equation 8).

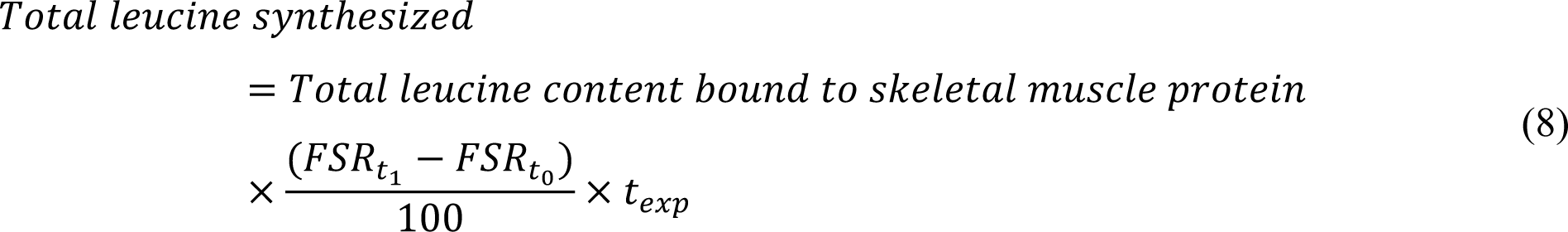

where *Total leucine content bound to skeletal muscle protein* is calculated as previously described, *FSR*_*t*0_ and *FSR*_*t*0_ are the FSR values at the start and end of the experimental intervention, respectively, in units of %/hour, and *t_exp_* is the experimental intervention duration in hours.

The phospho-protein data used in model calibration were measured using immunoblot analyses and were presented as fold changes from baseline, such that we converted the values to quantitative data that we could use in the model optimizer (Equation 9).

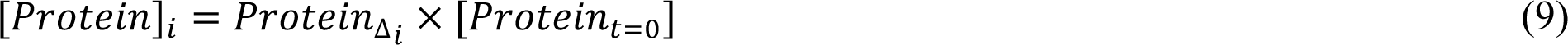

where [*Protein*]_*i*_ is the concentration of the protein at time *i*, *Protein*_Δ*i*_ is the fold change of the protein at time *i* determined by the immunoblot analysis, and [*Protein*]_*t*=0_ is the initial concentration of the protein.

The cost function for parameter optimization was the root mean square formula^63, 78, 79^ (Equation 10).

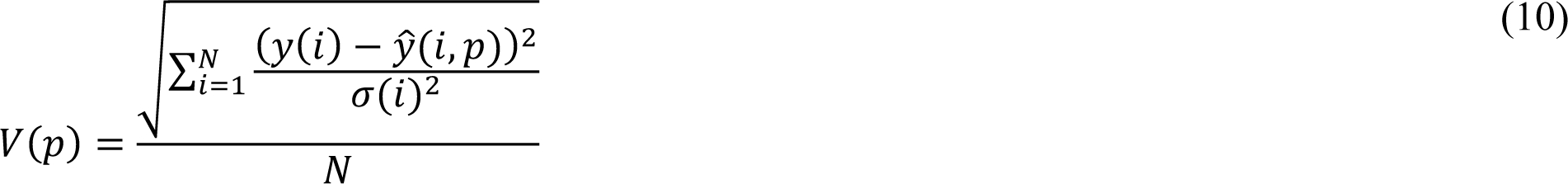

where *y*(*i*) represents the ith experimental data point, 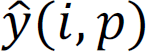 is the model predicted value given parameter *p*, σ(*i*) is the standard deviation of the experimental data, and *N* is the number of experimental data points for that parameter. The index *i* includes all time points at which each protein was measured.

### Model validation

We validated the calibrated model against six independent data sets with distinct feeding protocols: pulsatile feeding (i.e., four 0.9-gram boluses of leucine administered at 0, 45, 90, and 135 minutes^80^) and single boluses of leucine of different amounts (3.59 g^80^, 3.5 g^73^, 3.42 g^12^, 1.85 g^65^, and 1.8 g^81^; Table S7). We simulated the protocols by only altering the amount and timing of the ingested leucine to match the corresponding study protocols. No changes were made to any other model parameters or initial conditions in the model validation analyses.

### Calculations, numerical methods, and software

Net muscle protein balance (“net balance”, NB) was calculated as the difference between MPS and MPB (NB = MPS – MPB). Values of NB above zero indicate net synthesis while those below zero indicate net breakdown. CellDesigner 4.4 was used to illustrate all model topologies. R (version 4.2.1) was used to calculate the spline regression and to predict immunoblot-specific data. MATLAB version R2022a (9.12.0.2170939) was used for all model simulations, estimation of parameters, analyses, and calculations. We used the MATLAB ‘ode23s’ function to numerically integrate the model using default tolerances. The model code and data are available as a MATLAB package at https://github.sfu.ca/tmccoll/MuscleProteinSynthesisKineticModel/.

## Results

### Model development history

The development of our kinetic model of leucine-mediated signaling and protein synthesis was an iterative process wherein we added modules, adjusted reactions, meta-analyzed data sets, and added or removed molecules to arrive at a comprehensive and best-fit model. Essential steps of the model development history are described in the Supplementary Methods.

### Model calibration

We calibrated the model parameters in two steps: 1) the parameters from the insulin secretion module were independently fit to the insulin time-course (k51-67; volume-specific parameters were not adjusted), 2) the remaining model parameters were fit to all eight experimental datasets. Calibration of the model parameters resulted in a root mean square value of 5.66. The model simulations qualitatively agreed with experimental data for plasma leucine, intracellular leucine, plasma KIC, plasma insulin, phospho-Akt^S473^, phospho-p70S6K^T389^, and leucine incorporation to protein in response to a 3.5-gram bolus of leucine (Figure 2A).

**Figure 2.**
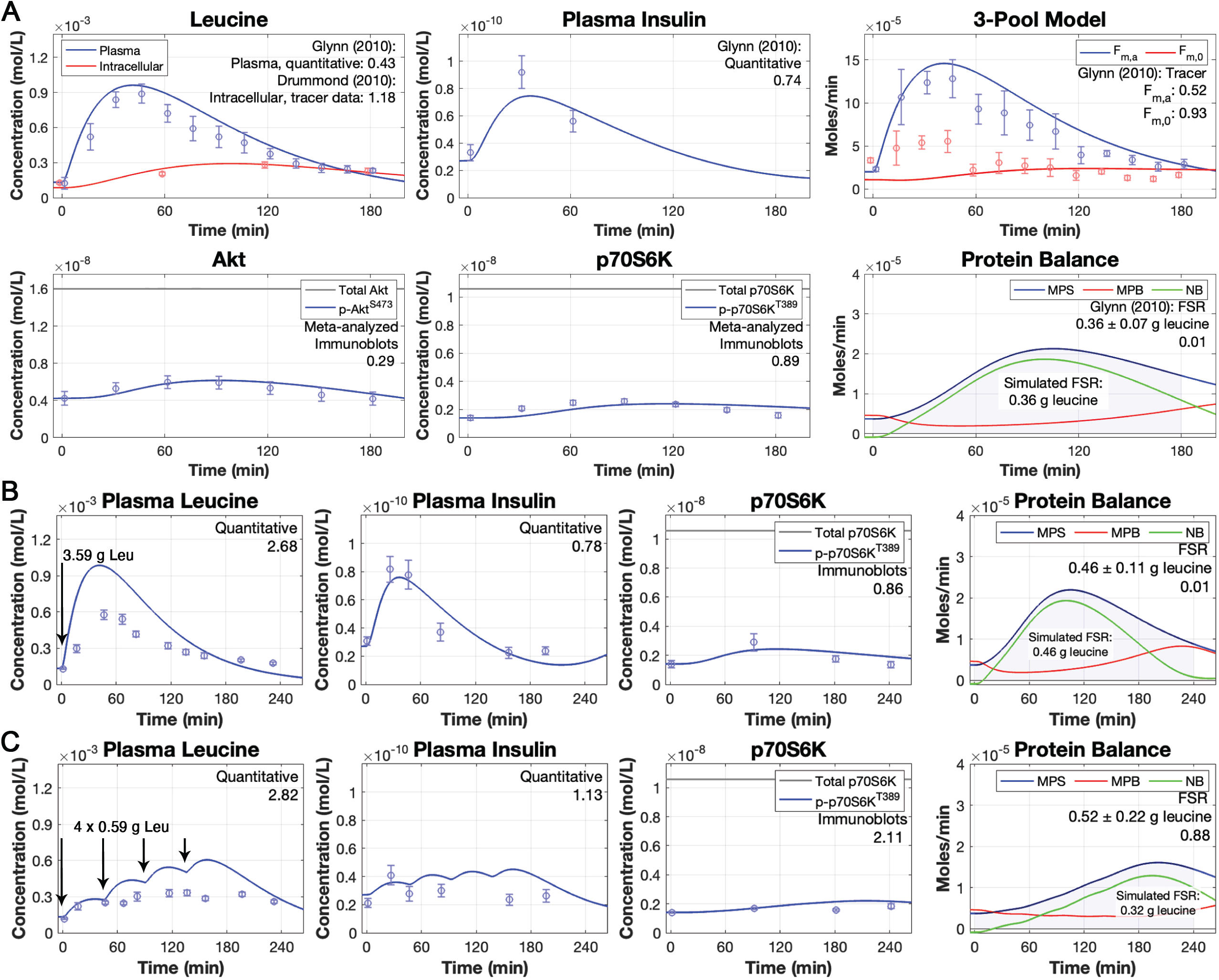
Model calibration and validation. (A) Simulated time-courses following model calibration of plasma leucine, intracellular leucine, plasma insulin, the three-pool model parameters F_m,a_ and F_m,0_, Akt (total and phosphorylated), p70S6K (total and phosphorylated), and muscle protein balance following a 3.5-gram bolus of leucine. Data points represent experimental data collected from two studies following the ingestion of a 3.5-gram bolus of leucine in human subjects^65, 73^. The data points for phospho-Akt^S473^ and phopsho-p70S6K^T389^ were predicted from spline regression equations obtained from meta-analyzed data. (B,C) Simulated time-courses of plasma leucine, plasma insulin, p70S6K, and muscle protein balance following either (B) a single 3.59-gram bolus of leucine^80^ or (C) pulsatile leucine feedings (0.59-grams of leucine provided at 0, 45, 90, and 135 minutes)^80^. Data points represent experimental data collected from the Mitchell et al.^80^ single bolus intervention (B) or pulsatile feeding intervention (C). Root mean square values for each time-course are included within each plot. The measured data are presented as means ± SE.

### Model validation

We comprehensively validated the model by comparing the output of the calibrated model against data from one pulsatile feeding protocol^80^ and five bolus feeding protocols^12, 65, 73, 80, 81^. The pulsatile feeding protocol was simulated by providing four 0.9-gram doses of leucine at 45-minute intervals, whereas the bolus feeding protocols were simulated by providing a 1.85-gram, 3,42-gram, 3.5-gram, or 3.59-gram bolus of leucine at 0 minutes. The pulsatile feeding protocol was distinct from the bolus feeding protocol that gave rise to the calibration data, such that it served as a stringent test for model validation. The model predictions for all six validation data sets qualitatively agreed with plasma leucine, intracellular leucine, plasma insulin, phospho-Akt^S473^, phospho-p70S6K^T389^, and MPS (Figure 2B,C; Figure S1).

### The model reveals discrepant experimental measurements

Our model highlighted discrepancies in experimental time-course data following leucine ingestion, in particular those pertaining to phospho-p70S6K^T389^ and plasma leucine. As previously discussed in the Methods, substantial differences existed in the estimates of phospho-p70S6K^T389^ dynamics from different lab groups. Figures S1A,F and the experimentally measured data from Glynn et al.^65^ (Figure B, Supplementary Methods, comprehensive model development procedure) show a large increase in phospho-p70S6K^T389^ at 60 minutes following leucine ingestion (39.5-, 12.5-, and 14.2-fold increase, respectively) that the model was unable to replicate. Comparatively, Figures S1B-D show a lower peak in phospho-p70S6K^T389^ (2.1-, 1.3-, and 1.6-fold increase, respectively) that the model accurately replicated. We searched for possible methodological differences or covariates to explain the discrepancies between the datasets but were unable to find any.

Discrepancies were also observed in the plasma leucine time-courses. We simulated the model against nine experimentally measured time-courses, in which the only change introduced to the model was the amount of leucine provided at time 0, which corresponded to the dose provided in the associated experimental data set. The model accurately simulated the plasma leucine time course in the calibration data set^65^ and in three validation datasets^65, 73, 81^ (Figure S2A-D). Each of these four interventions involved participants consuming a 10-gram bolus of EAA but with varying leucine content (1.8-to 3.5-grams). In contrast, the model overestimated the plasma leucine levels when leucine was administered in a larger, 15-gram EAA bolus^80^ (Figure S2E) and when leucine was administered in isolation^12^ (Figure S2F). When whey protein^75, 82^ or egg protein^83^ was used as the feeding intervention, the model over-estimated the plasma leucine levels and predicted an earlier peak (Figure S2G-I), thus suggesting the digestion and absorption kinetics are modulated when whey or egg protein is ingested compared to EAA alone. The amino acid compositions for each of the EAA interventions are listed in Table S8.

### Model analysis: simulation of unobserved variables

With the validated model 2, we analyzed the dynamics of model variables that are typically unmeasured in experimental studies (Figure 2: protein balance panels, Figure S3). For example, mTORC1 activity is difficult to measure experimentally. The mTORC1 kinase is regulated by the small GTPase Ras homolog enriched in brain (Rheb)^84^. Following leucine feeding, Rheb dissociates from TSC (TSC-Rheb, inhibitory state) allowing for mTORC1-Rheb co-localization, thereby increasing mTORC1 activity^84^. Quantification of the mTORC1-Rheb colocalization requires assays that measure protein proximity, such as immunofluorescence^85^, and these methods have seldom been employed to date in skeletal muscle metabolism research. In addition, the phosphorylation of the Ser2448 residue has been commonly but incorrectly measured as a proxy for mTORC1 kinase activity^61^. The model can simulate mTORC1 activity following leucine feeding and shows that little change in mTORC1 activity is required to induce the observed changes in MPS (Figure S3).

MPS and overall muscle protein dynamics are commonly inferred by the FSR in wet-lab experiments. However, FSR only accounts for MPS but not MPB, such that using FSR alone to infer the total muscle protein balance is predicated on the assumption that little to no change in MPB occurs following feeding. The model simulates MPS, MPB, and net balance following leucine intake, thus obviating the need to assume that MPS accounts for all changes in muscle protein balance. The model confirms that the MPS response following leucine feeding exceeds that of MPB, but that MPB is dynamic following feeding. Specifically, MPB decreases immediately after feeding in response to increased insulin concentrations, but then increases in the hours following feeding when insulin concentrations return to basal levels (Figure 2: protein balance panel). Prior to feeding, NB was negative, indicating net muscle protein breakdown. Post feeding, the combination of increased MPS and decreased MPB causes NB to be positive for several hours following feeding, corresponding to net muscle protein synthesis (Figure 2, protein balance panels).

### Model analysis: Knockdown of leucine signaling impairs muscle protein metabolism

Knockdown of leucine-mediated mTORC1 signaling caused a substantial loss of MPS (Figure 3). We simulated the knockdown of leucine-mediated mTORC1 activity by reducing the rate parameter controlling leucine-mediated mTORC1 activation (k39) by 1×, 0.75×, 0.50×, 0.25×, and 0.10× its calibrated value. All other model parameters remained the same. These adjusted models were then simulated using a single 3.5-gram bolus of leucine as input. Knockdown of mTORC1 activity reduced the downstream phosphorylation of p-p70S6K^T389^, which resulted in a proportional loss of MPS. The knockdown of mTORC1 activity did not affect MPB, such that there was a reduced net balance that corresponded to the loss in MPS.

**Figure 3.**
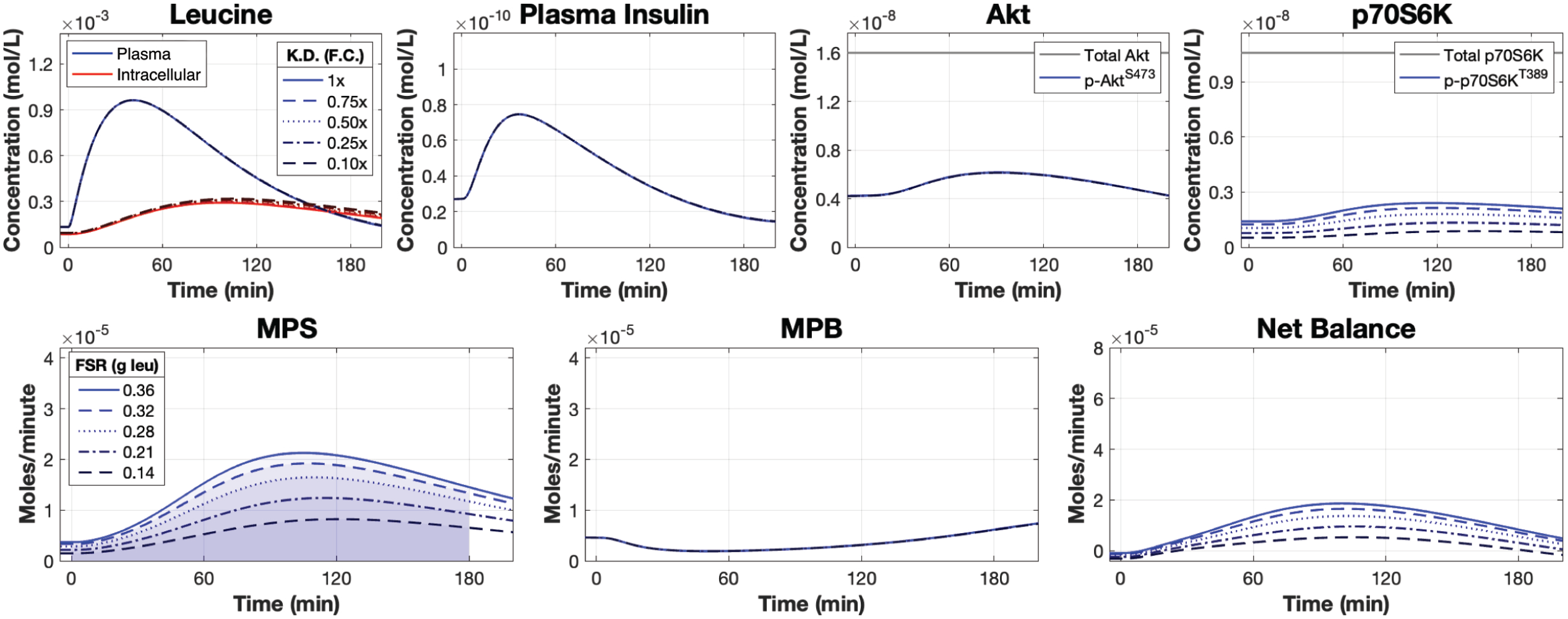
Knockdown of leucine signaling impairs MPS and net balance. Simulated time courses of plasma leucine, intracellular leucine, plasma insulin, Akt, p70S6K, and muscle protein balance following a 3.5-gram bolus of leucine with varying amounts of knock down to the rate controlling leucine-mediated mTORC1 activity. Signaling knockdown was simulated by modulating the kinetic rate parameter controlling leucine-mediated mTORC1 activity by 1×, 0.75×, 0.50×, 0.25×, and 0.10× its calibrated value. F.C. = fold change, K.D. = knockdown.

Knocking down leucine-mediated mTORC1 activity mimics the effects of rapamycin, a specific mTORC1 inhibitor in animals^13^ and humans^81^, which inhibits the early amino acid- and resistance training-induced increases in protein synthesis in humans^86^. To help validate our simulations and show the model’s potential usefulness for exploring rapamycin pharmacodynamics, we tested the model against the data from Dickinson et al.^81^. We simulated the model following the ingestion of a 1.8-gram bolus of leucine while progressively knocking down the rate controlling leucine-mediated mTORC1 activity (k39) until we obtained a qualitative match to the Dickinson et al.^81^ data set. We found that knocking down the rate controlling leucine-mediated mTORC1 activity 0.95× accurately simulated the loss of MPS and p70S6K signaling following leucine ingestion with rapamycin treatment (Figure S4)f. The loss of MPS was mediated by the loss of both basal and leucine-mediated phospho-p70S6K activity.

### Model analysis: Basal and activated leucine-mediated signaling contribute to controlling muscle protein metabolism

In analyzing the model, we noticed that changes to basal p70S6K values (i.e., both non-phosphorylated and phosphorylated p70S6K) influenced MPS. We pursued this observation by increasing the basal p70S6K values by a factor of two (2×) and four (4×) and simulated the model following a 3.5-gram bolus of leucine with all other parameters unchanged. When the p70S6K values were increased by a factor of two or four, MPS respectively increased 139% or 186% (Figure 4). The increase in basal p70S6K concentrations produced a marked increase in total p-p70S6K^T389^ signaling as measured by the AUC (1×: 3.72 × 10^-7^ mol×min/L, 2×: 5.52 × 10^-7^ mol×min/L, 4×: 8.01 × 10^-7^ mol×min/L). The change in p70S6K levels did not affect MPB, such that there was an increased net balance in both simulations that corresponded to the increase in MPS.

**Figure 4.**
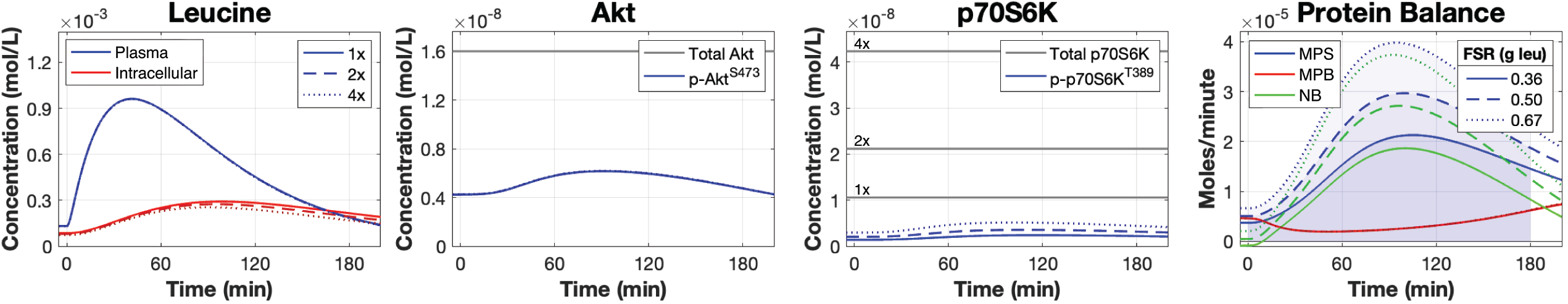
Increasing basal p70S6K levels substantially increases MPS. Simulated time courses of plasma leucine, intracellular leucine, Akt, p70S6K, and muscle protein balance following a 3.5-gram bolus of leucine with p70S6K concentrations (p70S6K and p-p70S6K^T389^) at the calibrated value (1×), two times the calibrated value (2×), and four times the calibrated value (4×).

Following the finding that basal p70S6K levels exert a strong influence on the MPS response, we tested whether leucine-mediated signal activation was necessary to produce a full MPS response when p70S6K levels were maintained at basal levels. The model was simulated with p70S6K levels maintained at their burn-in values and with mTORC1-mediated p70S6K phosphorylation inhibited following feeding. A 33% reduction in the MPS response was observed when mTORC1-mediated p70S6K phosphorylation was inhibited, with 0.24 g of leucine incorporated in the phospho-p70S6K-inhibited condition, compared to 0.36 g of leucine incorporated in the phospho-p70S6K responsive condition (Figure S5)

These results suggested that MPS is determined by both basal phospho-p70S6K levels and by mTORC1-mediated increased phosphorylation of p70S6K following feeding. The potential thus exists for one to compensate for the other. Indeed, reducing basal p70S6K levels by factors of 0.5 (0.5×), 0.25 (0.25×), or 0.125 (0.125×) caused respective reductions in MPS by 31%, 56%, and 72% (1×: 0.36 g leucine, 0.5×: 0.25 g leucine, 0.25×: 0.16 g leucine, 0.125×: 0.10 g leucine; Figure 5). However, independently increasing the rate controlling mTORC1-mediated p70S6K phosphorylation (k40) by 4× and 32× in the 0.5×- and 0.25×-p70S6K-level simulations, respectively, restored the MPS response (1× p70S6K, 1× mTORC1 kinase rate: 0.36 g leucine; 0.5× p70S6K, 4× mTORC1 kinase rate: 0.40 g leucine; 0.25× p70S6K, 32× mTORC1 kinase rate: 0.38 g leucine; Figure 5B,C). However, the MPS response was not restored in the 0.125×-p70S6K-level simulation even when mTORC1 kinase activity was increased by 64×, which produced a nearly saturated p-p70S6K^T389^ response (1× p70S6K, 1× mTORC1 kinase rate: 0.36 g leucine; 0.125× p70S6K, 64× mTORC1 kinase rate: 0.22 g leucine; Figure 5D). As with the previous simulations, no changes in MPB occurred with changes to the p70S6K levels or mTORC1 kinase activity, such that the changes in net balance corresponded exclusively to the changes in MPS.

**Figure 5.**
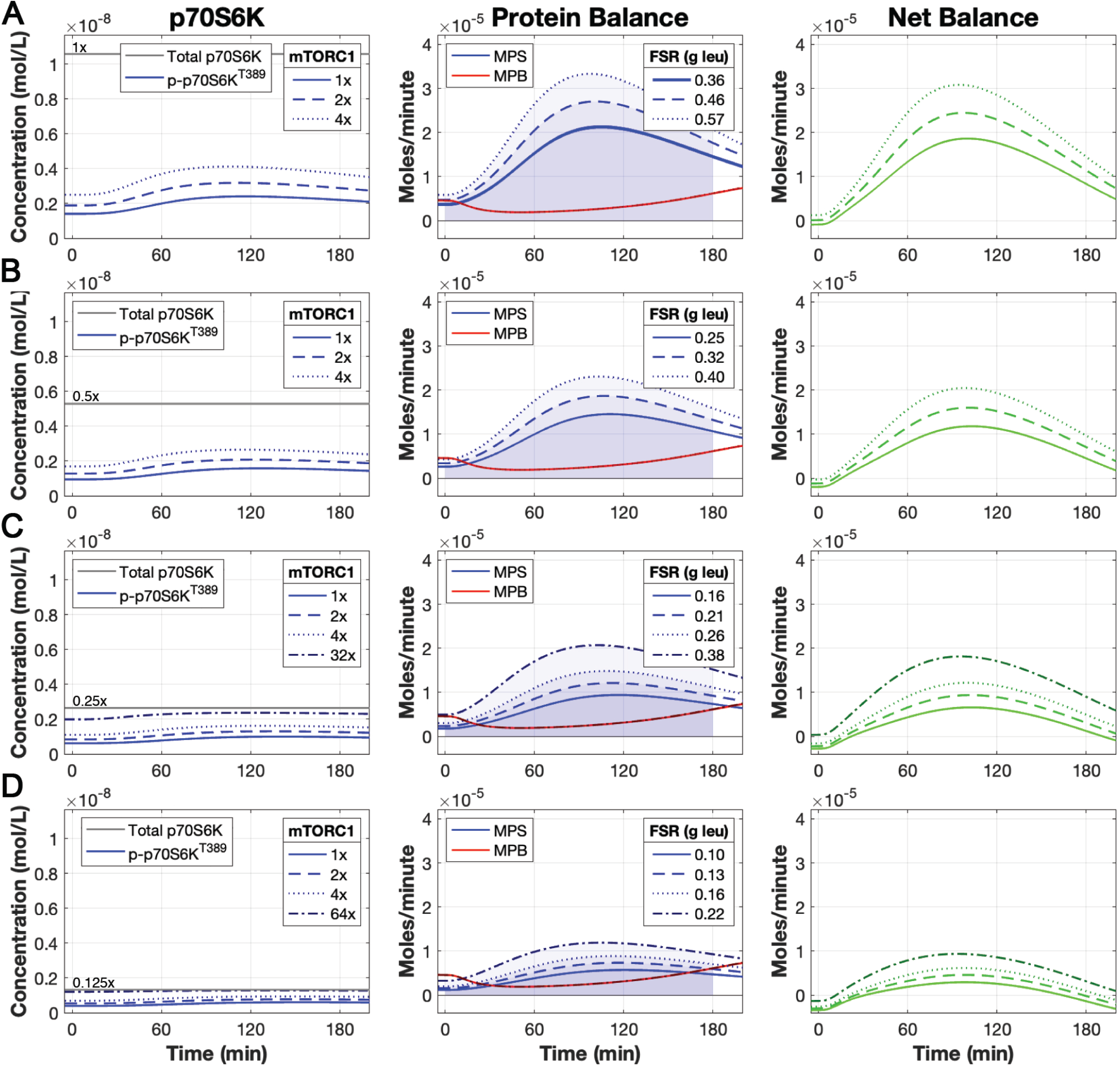
Enhanced signal activation can compensate for moderate losses in p70S6K levels. Simulated time courses of p70S6K, MPS, MPB, and NB following a 3.5-gram bolus of leucine with p70S6K concentrations (p70S6K + p-p70S6K^T389^) at (A) the calibrated value (1×), (B) 0.5-times the calibrated value (0.5×), (C) 0.25-times the calibrated value (0.25×), and (D) 0.125-times the calibrated value (0.125×). At each level of p70S6K, the rate controlling the mTORC1 kinase was simulated at 1×, 2×, and 4× of its calibrated value. The 0.25× and 0.125× p70S6K concentration were additionally simulated with the rate controlling mTORC1 kinase set at 32× and 64×, respectively. The baseline MPS time-course (1× p70S6K, 1× mTORC1 kinase) in (A) was bolded to serve as a reference line for (B), (C), and (D).

### Model analysis: The roles of insulin signaling in controlling muscle protein metabolism

Leucine and insulin both positively regulate mTORC1 activity and downstream MPS^18^, however, the contribution of insulin signaling to leucine-mediated MPS is unclear. Some studies suggest that insulin is required to induce a maximal MPS response^87, 88^, whereas others propose that it is not^34, 89^. To evaluate the contribution of insulin signaling on leucine-mediated MPS, we knocked down insulin signaling in the model in varying amounts (1×, 0.75×, 0.50×, 0.25×, 0.10×) and compared the resulting changes in muscle protein balance following the ingestion of a 3.5-gram leucine bolus. Specifically, we knocked down the rate controlling insulin-mediated IR_ý_ phosphorylation (k16) while keeping all other model parameters unchanged. We observed reduced Akt phosphorylation but no corresponding decreases in p70S6K phosphorylation or MPS (Figure 6). In fact, the model predicted a slight increase in MPS when insulin signaling was knocked down. However, the loss of insulin signaling resulted in a substantial increase in MPB since there was a loss of phospho-IR_ý_ mediated inhibition of MPB. Consequently, reduced net balance resulted from reduced insulin signaling because the increase in MPB surpassed the relatively small increase in MPS.

**Figure 6.**
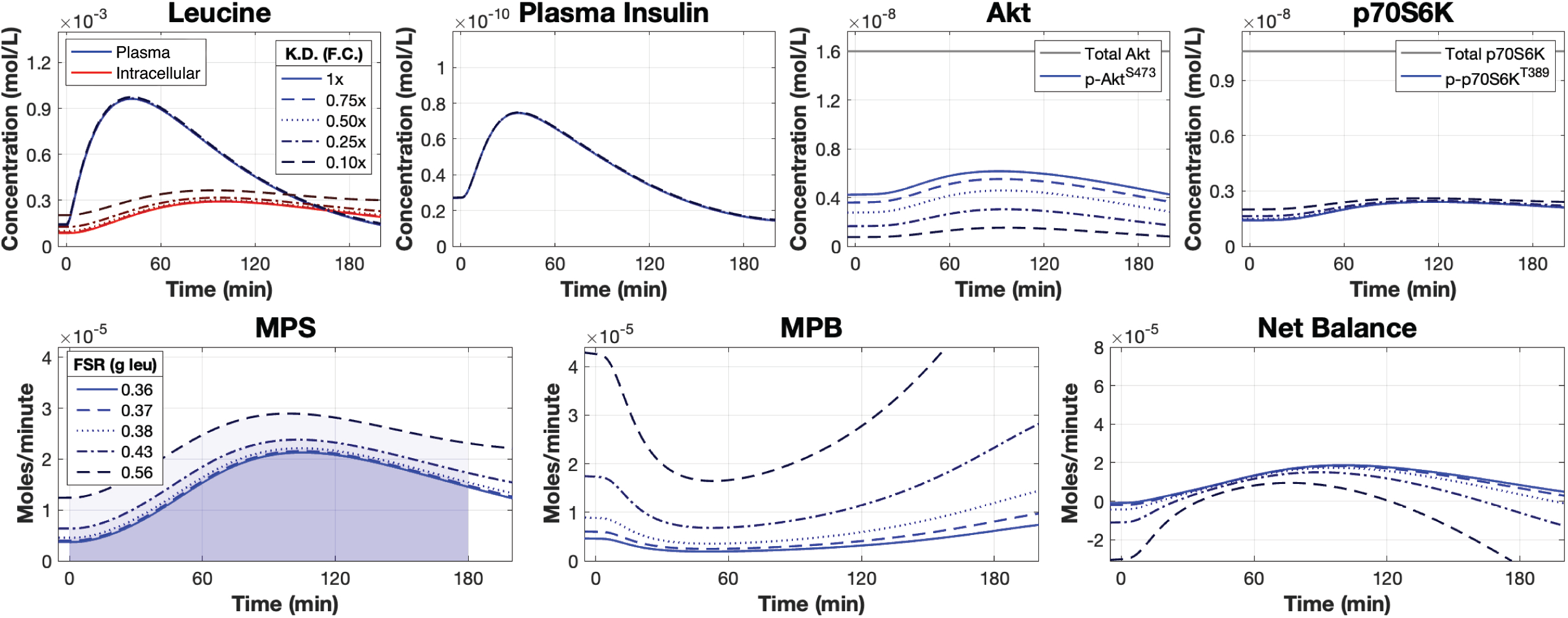
Knockdown of insulin signaling reduces net protein balance. Simulated time courses of plasma leucine, intracellular leucine, plasma insulin, Akt, p70S6K, and muscle protein balance following a 3.5-gram bolus of leucine with varying amounts of knock down to the rate controlling insulin-mediated insulin receptor (IR_ý_) phosphorylation. Signaling knockdown was simulated by modulating the kinetic rate parameter by 1×, 0.75×, 0.50×, 0.25×, and 0.10× its calibrated value. F.C. = fold change, K.D. = knockdown.

### Model analysis: The contributions of the p70S6K feedback pathways on muscle protein balance

Phospho-p70S6K^T389^ participates in two negative feedback pathways: one that promotes the phosphorylation of the serine residue on IRS1 and another that acts on mTORC1 to impair its kinase activity. We investigated the contributions of these feedback pathways to muscle protein balance. Specifically, we increased and decreased the kinetic parameters controlling p-p70S6K^T389^–mediated phosphorylation of the IRS1 serine residue (k21) and p-p70S6K^T389^– mediated inactivation of mTORC1 (k43) by 0.2×, 0.5×, 1×, 2×, and 5× and assessed the changes in muscle protein balance following the ingestion of a 3.5-gram leucine bolus. We found that increasing or decreasing k21 produced reciprocal changes in p-IRS1^S^, but muscle protein balance remained unchanged (Figure 7A). In contrast, adjusting k43 caused meaningful changes to mTORC1 activity and MPS. Specifically, increasing the strength of negative feedback on mTORC1 slightly reduced MPS while decreasing it led to increased MPS (Figure 7B).

**Figure 7.**
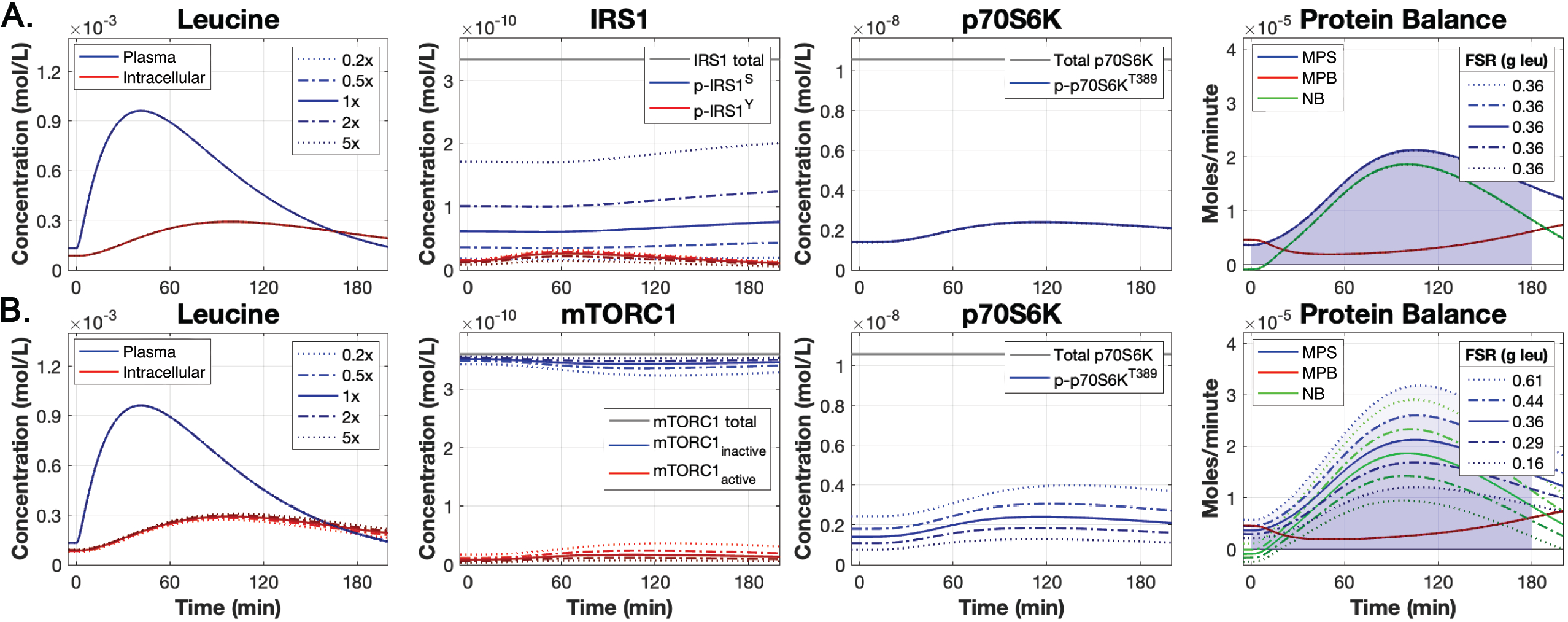
Contributions of phospho-p70S6K^T389^–mediated negative feedback on muscle protein balance. Simulated time courses following a 3.5-gram bolus of leucine with the kinetic parameters controlling (A) p-p70S6K^T389^–mediated phosphorylation of the IRS1 serine residue (k21) and (B) p-p70S6K^T389^–mediated inhibition of mTORC1 activity (k43) simulated at 0.2×, 0.5×, 1×, 2×, and 5× their calibrated value. Total IRS1 includes the PI3K p-IRS1^Y^ complex and the non-phosphorylated IRS1 protein, neither of which were presented in the IRS1 plot.

## Discussion

Skeletal muscle metabolism is complex and involves the dynamic interplay of multiple cellular and whole-body physiological processes. Experimental studies have examined these processes in relative isolation, such that how these processes operate as a system remains poorly understood. In this study, we developed and analyzed a kinetic model of protein translational signaling and protein metabolism in human skeletal muscle cells in response to leucine ingestion. Our primary objective was to create a first-generation model of the signaling controlling protein metabolism in human skeletal muscle that incorporated physiologically realistic input dynamics to drive the downstream protein signaling and metabolic responses. We accomplished this goal by modifying and amalgamating published models of mTOR signaling^46^, skeletal-muscle leucine kinetics^43^, and insulin dynamics^41^ and then updating the overall model topology according to the latest literature. The resulting model satisfactorily predicts data collected from human participants in response to various leucine feeding interventions. Our model revealed three key findings: 1) basal levels of p70S6K are an important determinant of MPS rates, the contribution of which was greater than increased p70S6K phosphorylation status alone; 2) insulin signaling can strongly influence muscle protein balance through its inhibition of MPB; and 3) p70S6K-mediated negative feedback of mTORC1 signaling reduces MPS in a dose-dependent manner. Our study thus motivates new hypotheses regarding the mechanism by which signaling influences protein metabolism, reconciles controversial aspects of protein metabolism, and provides a foundation for future modeling studies of skeletal muscle protein metabolism.

### Main findings and their implications

A key finding from our model analysis was that basal levels of p70S6K substantially influenced MPS. Most experimental studies assess the relative differences in phospho-p70S6K levels between interventions and use these data to infer MPS. Although phospho-p70S6K controls MPS^31, 81^ and our model confirms the need for phospho-p70S6K signaling for MPS (Figure 5, Figure S5), our results suggest that differences in basal levels of p70S6K may have a more pronounced influence on MPS than the enhancement caused by leucine-stimulated mTORC1 signaling. Of the experimental studies we found that featured amino-acid or whey protein feedings, none quantified the absolute concentrations of signaling proteins (phosphorylated or non-phosphorylated).

This result has important health implications because differences in basal p70S6K levels may determine an individual’s susceptibility to sarcopenia and responsiveness to resistance training. Sarcopenia is thought to be primarily due to *anabolic resistance* (reduced MPS response) to feeding^22^. Our results suggest that a cause of anabolic resistance may be reduced p70S6K protein levels. Indeed, Cuthbertson et al (2005)^90^ found that the total concentration of p70S6K protein in elderly men was 50% of that in young men, and this corresponded to ∼30-40% less myofibrillar FSR in the elderly men following EAA feeding. In addition, our model suggests that modest losses in p70S6K protein levels can be compensated for by increased mTORC1 kinase activity. This scenario appears to be supported from two experimental studies. Markofski et al.^91^ found an increase in basal p70S6K^T389^ phosphorylation in the older adults group but no difference in basal FSR between young and older adults. Cuthbertson et al.^90^ observed a statistically non-significant increase in basal p70S6K^T389^ phosphorylation in the elderly group in the basal condition and in response to low-dose EAA feedings (10 grams). The increased basal p70S6K^T389^ phosphorylation potentially allowed for the similar MPS responses between the old and young groups. By comparison, resistance training induces an anabolic state wherein MPS is elevated beyond MPB to allow for an increase in net balance and muscle mass^22^. Studies in rats show that the levels of several mTOR signaling proteins, including p70S6K, may be increased following chronic resistance training when biopsies are extracted following the recovery from exercise (i.e., 24-28 hours post exercise bout)^92, 93^. We postulate that the increase in these mTOR signaling proteins may contribute to the increase in MPS and the resulting increase in muscle mass.

Overall, these results suggest that an individual’s anabolic signaling system can mechanistically act in one of two ways to support MPS: 1) a “permissive” mechanism or 2) an “activation-dependent” mechanism. In the “permissive” mechanism, relatively high basal levels of phospho-p70S6K are sufficient to support MPS when amino acid availability increases. Leucine-mediated activation of mTORC1 contributes to MPS but to a relatively minor extent. In this case, the system is always “primed” for when amino acids are present. In the “activation-dependent” mechanism, basal phospho-p70S6K levels are relatively low and insufficient to support adequate MPS without additional leucine-stimulated mTORC1 activation.

An important benefit of our model is that it enables the simulation of all species within the system, which enables investigation of species that are commonly overlooked in experimental interventions. For example, MPS, MPB, and muscle protein balance (i.e., NB) are all simulated. The literature often states that MPS is more sensitive to anabolic stimuli compared to MPB and is the primary determinant of changes to muscle protein balance^21, 94^. Additionally, MPB is more methodologically challenging to measure compared to MPS^95^. For these reasons, experimental studies commonly feature MPS (i.e., FSR) only, which is used to infer changes to muscle protein balance. However, ignoring MPB will cause one to overlook its contributions to muscle protein balance and may cause misinterpretations regarding the role of insulin. Insulin signaling has two primary roles in the system: 1) it stimulates mTORC1 activity and 2) it inhibits MPB. Controversy exists regarding the influence of insulin on muscle anabolism. Studies in rats document reduced MPS when insulin secretion was blocked^88^ or maintained at basal levels^87^ following amino acid feeding. In contrast, studies in humans report unchanged MPS in response to casein protein ingestion^89^ or intravenously administered amino acids^34^ alongside insulin clamped at higher-than-systemic postabsorptive concentrations. We sought to assess the role of insulin on muscle anabolism by knocking down insulin signaling in the model. We found that knocking down insulin signaling had little effect on MPS but resulted in substantially increased MPB, which in turn reduced muscle protein balance. Therefore, studies in which only MPS was measured may conclude that insulin has little influence on muscle anabolism, whereas considering both MPS and MPB leads to the hypothesis that the increase in MPB following insulin knockdown reduces muscle protein balance to a quantitatively important extent.

We also assessed two p70S6K-mediated negative feedback pathways to better understand their control over muscle protein metabolism. One pathway features p70S6K-mediated phosphorylation of the serine residues of IRS1^28, 29, 31^. Serine-phosphorylated IRS1 promotes the degradation of IRS1, thereby negatively regulating insulin signaling^31^. Our analysis of this pathway confirmed that p70S6K controlled the level of serine phosphorylated IRS1 and exerts slight control of downstream signaling (i.e., p-Akt^S473^; data not shown), but this effect did not propagate to meaningful changes in MPS. This result is consistent with our previous result that insulin-knockdown had little effect on MPS, and with other studies that showed that insulin signaling contributes little to MPS^34, 89, 90^. The other feedback pathway was p70S6K-mediated phosphorylation of the Ser2448 residue on mTOR^96^. This residue resides in the mTOR negative regulatory domain^61^. Phosphorylation of the Ser2448 residue reduces mTORC1 kinase activity^61, 96–98^, thereby reducing MPS. Our analysis of this feedback loop showed a dose-response relationship with MPS. A poorly understood phenomenon in muscle protein metabolism is the “muscle-full effect,” in which MPS decreases post feeding despite continually elevated plasma and intracellular EAA and leucine levels^75, 99^. We wonder whether the p70S6K/mTORC1 negative feedback loop could contribute to the muscle-full effect^75^. This hypothesis could be experimentally tested by inhibiting the p70S6K/mTORC1 negative feedback pathway by blocking the Ser2448 binding site or removing the residues in the mTOR negative regulatory domain, and measuring MPS in response to EAA or whey protein feeding.

### Validity of the results

The robustness of the above findings is ultimately predicated on the validity of the model, which we took great lengths to establish. The model calibration process integrated several data types, including quantitative data (e.g., plasma leucine, plasma insulin), isotopic tracer data (e.g., intracellular leucine, 3-pool parameters, FSR), and semi-quantitative immunoblots (e.g., phospho-Akt^S473^, phospho-p70S6K^T389^). Due to the semi-quantitative nature of immunoblot data and the variability in immunoblot measurements^100^, we incorporated a meta-analytic approach to attempt to increase the accuracies of the phospho-Akt^S473^ and phospho-p70S6K^T389^ time-courses (described in the Supplementary Methods). We used the regression model from the meta-analysis to impute time-course data for each species in 30-minute increments (i.e., six data points over the 180-minute calibration period), which provided time courses of higher resolution for data fitting compared to single experimental time courses. Additionally, we used selected baseline 3-pool model parameter values^39^ and the baseline KIC oxidation rate^44^ to calibrate the basal skeletal-muscle leucine module.

Once the model was satisfactorily calibrated, we searched extensively for appropriate validation data sets. We located six such data sets, five of which featured single, bolus feedings of varying leucine doses (1.85-to 3.59-grams), and one that featured a pulsatile feeding protocol, i.e., repeated small doses of leucine. The pulsatile feeding intervention served as a particularly stringent test for model validation. By adjusting only the timing and dose of leucine ingestion in the model (i.e., all other model parameters were unchanged) across these heterogenous interventions (i.e., each study featured variable participant characteristics, differences in ingested solutions, etc.), the model showed a remarkable ability to successfully predict qualitative features such as the timing of peak concentrations or rates and the overall dynamics of the variables measured in the six validation data sets. Several cases existed in which the model simulations differed from the measured data, but these discrepancies were typically slight over- or under-estimations and the model predictions still captured the trends in the data. The model showed a strong ability to fit phospho-data time-courses, except for two, but even in these cases the model achieved satisfactory error cost values (discussed below). Furthermore, the model achieved good fits for all intracellular leucine simulations and F_m,a_ simulations.

The model predicted four aspects of the validation data less well, but plausible explanations exist for most of these cases. First, the model overpredicted the plasma leucine concentrations relative to the measured values in three of the six datasets (Figure S1B-D). We propose that these discrepancies were in part due to the different nutrient formulations fed to the participants, the effects of which we discuss in detail in the following subsection. Second, in two of the bolus feeding validation data sets^65, 81^, the phospho-p70S6K^T389^ experimental measurements appeared to be overestimated and exceeded what the model could simulate. We propose that these discrepancies likely arose from methodological differences in data collection and analysis, discussed in detail in the Supplementary Methods. Third, in the Glynn et al.^65^ 1.85-gram leucine intervention, a high plasma insulin response at 30-minutes was observed that the model did not replicate. Leucine is an insulin secretagogue, such that a leucine dose-insulin response relationship should exist^23, 24^. However, the reported plasma insulin response was excessive compared to other interventions that provided greater leucine doses [3.42-grams^12^, 3.5-grams^65^, 3.59 grams^80^] but that reported lower plasma insulin levels at 30 minutes. Fourth, the model underestimated the total leucine synthesized over the intervention period (i.e., an extrapolation of FSR) in three of the validation datasets ^12, 80, 8180^. Indeed, substantial variability in experimentally measured leucine synthesis (i.e., FSR) was observed: Glynn et al.^65^ and Dickinson et al.^81^ applied similar feeding interventions (1.85 g leucine, 10 g EAA and 1.8 g leucine, 10 g EAA, respectively) but reported markedly different amounts of leucine synthesized, i.e., 0.11 ± 0.02 g at 180 minutes versus 0.32 ± 0.07 g at 120 minutes, respectively. Despite these instances of lack of fit, on balance the model accurately replicated the biological responses to various leucine feeding interventions.

### Impacts of modeling decisions

Any model must balance comprehensiveness with parsimony, and we discuss here four of our major decisions. First, we made the simplifying assumption to focus the model exclusively on leucine, despite leucine being just one of 20 amino acids that comprise human proteins. Leucine was the focus because it is the most important amino acid with respect to anabolic signaling^12–15^, and including other amino acids would have greatly increased the model’s complexity. The impacts of this decision cannot be definitively evaluated at present, but our results indicate that discrepancies may be introduced at the level of leucine digestion and absorption. Specifically, we compared our model to data from feeding interventions featuring differing nutrient compositions, including leucine alone, EAA solutions, and whey or egg protein. In the case of whey and egg protein, the model overestimated the plasma leucine response with earlier peak concentrations in each dataset (Figure S2G-I). This finding makes sense because both protein sources consist of whole proteins (whey is a component of whole milk), which require digestion unlike the free-form amino acids in EAA. Thus, leucine absorption is delayed when administered in whey protein^12, 101^. Simulation of more complex protein sources would require more details regarding protein digestion and absorption^102–104^, which represents a future direction for our model. In addition, the model was able to accurately simulate four plasma leucine time-courses following varying leucine boluses (1.80 to 3.50 grams) provided as part of a 10-gram EAA solution (Figure S2A-D). However, the model overestimated the plasma leucine response in two data sets, one that provided 3.59-grams of leucine with 15-grams of EAA^80^ (Figure S2E) and another that provided 3.42-grams of leucine alone^12^ (Figure S2F). In accordance with the whey protein results, the greater amount of EAA in the Mitchell study may have reduced the amount of leucine absorbed, thus causing the overprediction (Figure S2E). However, our model should have underpredicted the plasma leucine data of Wilkinson but instead it overpredicted them (Figure S2F). Collectively, these results indicate that different nutrient formulations may affect leucine absorption dynamics, but lab-specific differences in experimental methods and random variability may also affect plasma leucine data. Future versions of the model will examine the effects of the other feeding interventions.

Second, the model does not simulate the muscle-full effect. Maximal MPS rates occur following the ingestion of ∼10-grams of EAA^82, 90, 105, 106^ every 3 hours^105^; increasing the feeding dose or frequency does not further increase MPS. Therefore, the current model may overestimate MPS following large or frequent doses of leucine. Although this issue may be problematic when attempting to use the model to determine optimal feeding profiles, we do not believe that any interventions simulated within this study contained sufficiently high doses of leucine to elicit the muscle full effect, such that this limitation should not influence our findings.

Third, we made the simplifying assumption to use ODEs to encode the model, which requires the assumption that all biochemical reactions occur deterministically in a homogenous compartment and do not account for spatially distributed cellular processes. Regarding protein localization, both the insulin- and leucine-dependent pathways interact with proteins bound to the surface of lysosomes (e.g., Rheb, Ragulator) prior to activating mTORC1^26, 107^. To account for spatially localized species, we have assumed the model to be a well-mixed compartment and we have modelled any changes in species localization (i.e., movement between the blood plasma and cellular space) with ODEs through elementary reactions between different compartments, resolving the requirement to model changes in concentration with respect to space^40^.

The fourth decision we made was to retain most of the components of the mTOR signaling network, which made the model relatively complex. Complex models can be limited in several ways, such as challenges in estimating the parameter values, overfitting the data, and reduced tractability. Some may argue that a more parsimonious model would be better. We justify the model’s scope as follows. First, we emphasize that we did simplify parts of the model, in particular removing proteins that did not influence model dynamics (e.g., PRAS40). Second, many of the represented reactions depict “lumped” processes in which distinct molecular processes are collectively represented by one rate equation. For example, the mechanisms of leucine sensing are implicitly represented within reaction 39 (Table S2). Therefore, the model is relatively simple compared to biological reality. Third, the experimental data used for model calibration featured model components that were distributed throughout the model topology, including at the input (e.g., plasma leucine, insulin), leucine kinetics (e.g., intracellular leucine, 3-pool model), central signaling network (p-Akt^S473^), downstream signaling level (p70S6K^T389^), and final output (MPS). Therefore, we could determine whether the model was operating correctly, and if not, at what level the model failed to replicate the experimental data^108^. Furthermore, there was no evidence of model overfitting; rather the model still exhibits lack of fit in places (discussed above). Finally, the current form of the model provides the potential for enhanced mechanistic insights and facilitates future hypothesis generation regarding the mechanisms of disease states. For example, insulin resistance involves IRS1 phosphorylation modulated by protein tyrosine phosphatase 1B (PTP1B)^29^ and modulated feedback between p70S6K and IRS1^79^. In addition, the anabolic resistance of sarcopenia could involve any of the components in the signaling network, which our model is well positioned to explore.

### Practical implications

Our study represents the first model that simulates skeletal muscle protein metabolism in humans using realistic whole-body dynamics of leucine and insulin following feeding. Previous mathematical models of translational signaling and protein metabolism have been limited to *in vitro* cell lines (e.g., HeLa cells, C2C12 myoblasts, CHO cells) and used non-physiological input dynamics (e.g., constant inputs). Therefore, our model has the advantage of being directly applicable to human skeletal muscle. Furthermore, our model can simultaneously fit, and therefore reconcile, data from distinct experimental methods, e.g., liquid chromatography-based measures of plasma amino acid levels, ELISA-based measures of plasma hormone levels, immunoblotting of signaling proteins, and metabolite fluxes from stable isotope tracers. The integration of various data types into a single framework allows for a clear representation of the multi-scale system through which leucine functions and can act as a tool for understanding the complex components of skeletal muscle metabolism (e.g., 3-pool model, leucine kinetics). Findings from our model analysis suggest the need to quantify in absolute units total and phospho-protein levels of key signaling proteins (e.g., p70S6K) to better understand their role in MPS control. These differences in basal protein levels may contribute to differences in MPS rate, which could lead to the development of sarcopenia. Additionally, our model highlights the lack of phospho-protein time-course data in many proteins of the mTOR signaling network following feeding interventions. Although phospho-Akt and phospho-p70S6K can inform much of the insulin- and leucine-induced signaling activity, the measurement of additional phospho-proteins throughout the signaling network following human interventions would allow for a better understanding of the overall network dynamics.

### Conclusions

In summary, we have developed and analyzed a mathematical model of protein translational signaling in human skeletal muscle cells following leucine feeding that features hormonal and nutritional inputs. Our findings suggest that basal levels of p70S6K have a key influence on MPS rates that is greater than that of dynamic changes in phospho-p70S6K levels, that insulin signaling plays a prominent role in muscle protein balance through its effects on MPB, and that p70S6K-mediated feedback may restrict MPS. Our model provides an essential tool for integrating diverse data types, reconciling contradictory data, and for systematically investigating the various multi-level mechanisms governing skeletal muscle protein metabolism. The model may therefore be used to generate more informed and sophisticated hypotheses for experimental testing.

## Acknowledgements

We thank Timothy Rattan for his contribution in curating calibration and validation data, Han Jie Liu for his contribution in curating validation data, and Marvin K.F. Ly for his research pertaining to muscle protein breakdown and leucine-mediated mTORC1 activity.

This work was supported by a Natural Sciences and Engineering Research Council of Canada (NSERC) Collaborative Research and Training Experience scholarship to T.J.M. and a NSERC Discovery Grant to D.C.C. (RGPIN 06004-2014). The funders had no role in study design, data collection and analysis, decision to publish, or preparation of the manuscript.

## Author contributions

Conceptualization, T.J.M. and D.C.C.; Methodology, T.J.M and D.C.C.; Software, T.J.M.; Validation, T.J.M; Formal Analysis, T.J.M. and D.C.C.; Investigation, T.J.M. and D.C.C.; Resources, D.C.C.; Data Curation, T.J.M.; Writing – Original Draft, T.J.M and D.C.C.; Writing – Review & Editing, T.J.M and D.C.C.; Visualization, T.J.M.; Supervision, D.C.C.; Project Administration, D.C.C.; Funding Acquisition, D.C.C.

## Supplementary information titles and legends

**Figure S1. Related to Figure 2 Comprehensive model validation.**

Simulated time-courses of plasma leucine, plasma insulin, Akt, p70S6K, three-pool model parameters F_m,a_ and F_m,0_, and muscle protein balance following a single (A) 1.85-gram, (B) 3.59-gram, (D) 3.42-gram, (E) 3.50-gram, or (F) 1.80-gram bolus of leucine or (C) pulsatile leucine feedings (0.59-grams of leucine provided at 0, 45, 90, and 135 minutes). Data points represent experimental data collected from (A) Glynn et al.^65^, (B,C) Mitchell et al.^80^, (D) Wilkinson et al.^12^, (E) Drummond et al.^73^, and (F) Dickinson et al^81^. Root mean square values for each time-course with corresponding experimental data are included within the respective plot. The measured data are presented as means ± SE.

**Figure S2. Related to Figure 2. Discrepant plasma leucine measurements between different data sets.**

Simulated plasma leucine time-courses in response to bolus leucine feedings (A-F: 3.5 g, 1.85 g, 3.50 g, 1.80 g, 3.59 g, and 3.42 g, respectively) and whey protein feedings with differing leucine amounts (G-I: 4.1 g, 2.1 g, or 1.67 g, respectively). Data points represent experimental data from (A) Glynn et al.^65^, (B) Glynn et al.^65^, (C) Drummond et al.^73^, (D) Dickinson et al.^81^, (E) Mitchell et al.^80^, (F) Wilkinson et al.^12^, (G) Atherton et al.^75^, (H) Moore et al.^82^, and (I) Mazzulla et al.^83^. The ingested dose of leucine is listed on the left side of each plot and the root mean square values for each time-course is listed in the upper-right portion of each plot. The measured data are presented as means ± SE.

**Figure S3. Related to Figure 2. Simulation of unmeasured species in the model.**

Simulated time-courses of all model species that were not experimentally measured following a 3.5-gram bolus of leucine. The phospho-Akt^S473^ time-course includes the predicted time-course data from the meta-analyzed spline regression (mean ± SE). Total IRS1 includes the PI3K p-IRS1^Y^ complex, which is presented in the adjacent PI3K panel.

**Figure S4. Related to Figure 3. Model simulation of rapamycin ingestion.**

Simulated time-courses of plasma leucine, intracellular leucine, plasma insulin, p70S6K, and muscle protein balance following a 1.8-gram bolus of leucine with prior ingestion of rapamycin, a potent mTORC1 inhibitor. Data points represent experimental data from Dickinson et al.^81^. Root mean square values for each time-course with corresponding experimental data are included within each plot. The measured data are presented as means ± SE.

**Figure S5. Related to Figure 3. Phospho-p70S6K signaling is required to elicit a maximal MPS response.**

Simulated plasma leucine, intracellular leucine, p70S6K, and muscle protein balance time courses following a 3.5-gram bolus of leucine with (A) phospho-p70S6K sensitive to leucine-mediated signaling or (B) phospho-p70S6K insensitive to leucine-mediated signaling and maintained at the calibrated concentration over the simulation duration. Data points in (A) represent data from the model calibration. The gray dashed line in the (B) protein balance plot is a reference line for the original MPS from (A). Root mean square values for each time-course with corresponding experimental data are included within each plot. The measured data are presented as means ± SE.

**Table S1. Related to Figure 2.** Proteins included in the model and their cellular properties.

**Table S2. Related to Figure 2.** Reaction rate equations.

Reaction rate equations and the functional relationships used in the oscillatory insulin module.

**Table S3. Related to Figure 2**. System of ordinary differential equations.

**Table S4. Related to Figure 2**. Model species and initial concentrations.

Model species (state variables) and their initial concentrations before and after the model calibration and at the end of the model equilibrium period.

**Table S5. Related to Figure 2**. Model parameter values.

Model parameter values before and after model calibration.

**Table S6. Related to Figure 2A**. Calibration dataset characteristics.

**Table S7. Related to Figure 2B,C.** Validation dataset characteristics.

**Table S8. Related to Figure 2.** Amino acid profile for each intervention.

