## Supplementart methods for "Kinetic modeling of leucine-mediated signaling and protein metabolism in human skeletal muscle"

#### Contents:

Supplementary methods

*Model development: Version 1 of the model exhibits lack of fit to experimental data*

*Model development: Meta-analysis of phospho-p70S6K$^{T389}$ and phospho-pAkt$^{S473}$ time course data following leucine feeding*

*Model development: Including an insulin secretion module to simulate the oscillatory secretion of insulin*

### Supplementary methods

*Model development: Version 1 of the model exhibits lack of fit to experimental data*

Calibration of the model parameters in version 1 of the model (Supplementary Figure A) resulted in a root mean square value of 7.70. The model simulations qualitatively agreed with experimental data (i.e., it displayed similar timing of peak concentrations or rates and overall dynamics) for plasma leucine, intracellular leucine, $F_{m,a}$, $F_{m,0}$, phospho-Akt$^{S473}$, and MPS in response to a 3.5-gram bolus of leucine (Supplementary Figure B). An obvious lack of fit existed for phospho-p70S6K$^{T389}$, particularly at 60 minutes (Supplementary Figure B), which was not improved by manually adjusting the rate parameters that directly influenced phospho-p70S6K$^{T389}$ kinetics (e.g., k40-42). Additionally, there was poor fit for plasma insulin wherein there was a delayed peak concentration, and the peak insulin concentration did not reach that of the experimental data.

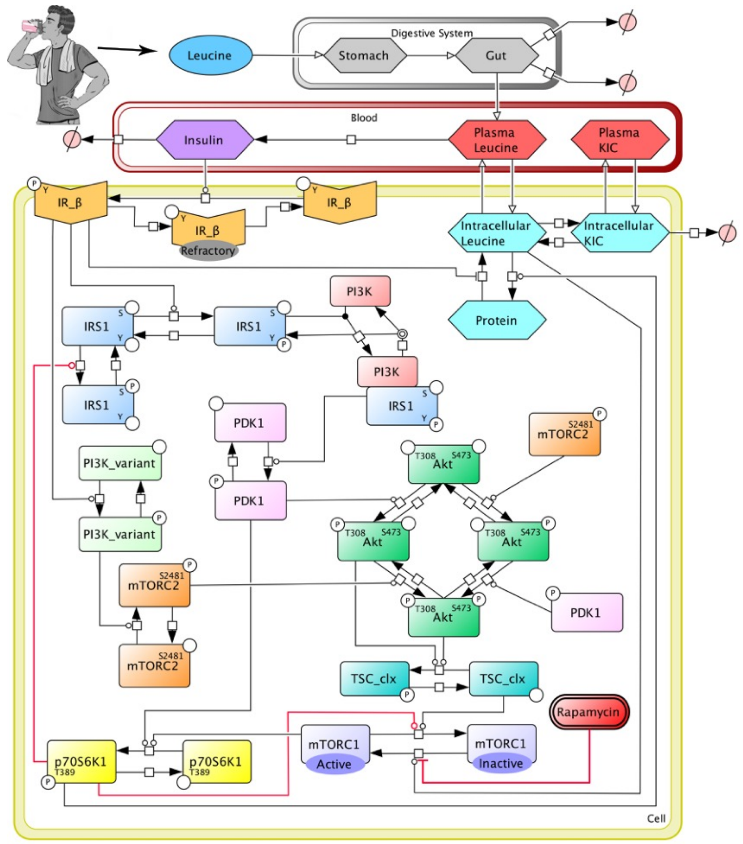


**Figure A. Topology of the version 1 model.** Model reaction diagram of protein translational signaling in response to leucine ingestion. mTORC2$^{S2481}$ and PDK1$^{P}$ are represented as independent species for the phosphorylation of Akt$^{S473}$ and Akt$^{S473,T308}$, respectively, but both are subject to the same control as the integrated species. ). Transfer of masses are denoted by open headed arrows. Chemical reactions are denoted by solid headed arrows. Inhibition is denoted by a flat-headed line. P = phospho residue, S = serine, T = threonine, Y = tyrosine.

*Model development: Meta-analysis of phospho-p70S6K$^{T389}$ and phospho-pAkt$^{S473}$ time course data following leucine feeding*

The inability of the model to fit the phospho-p70S6K$^{T389}$ time-course from Glynn et al.[1] prompted us to examine in detail the existing data regarding p70S6K phosphorylation following feeding. We

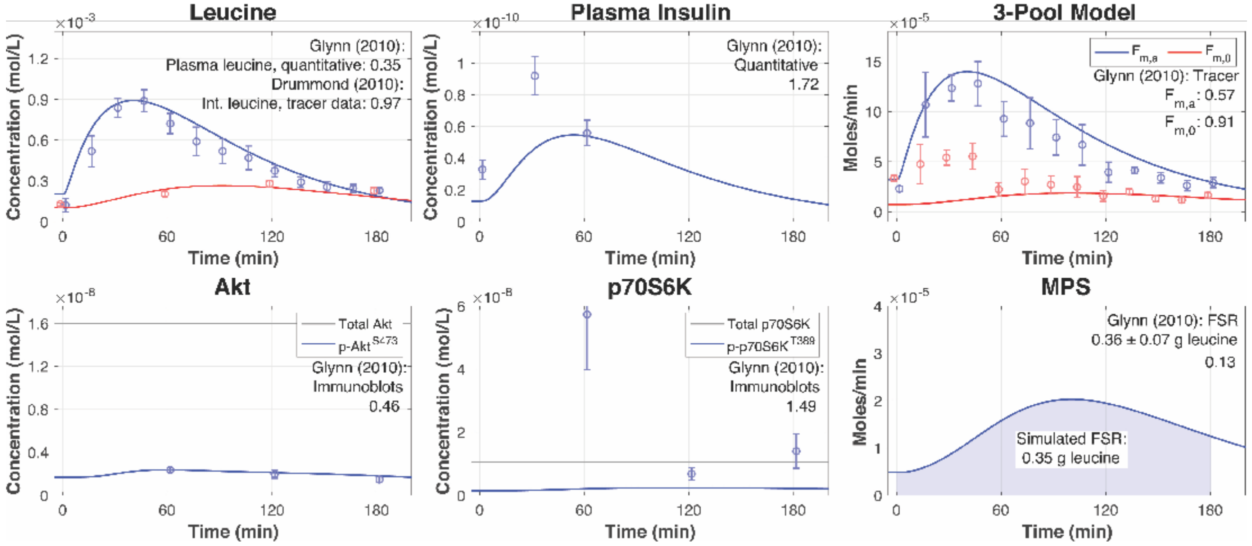


**Figure B. Calibration of model version 1.**

Simulated time-courses following model version 1 calibration of plasma leucine, intracellular leucine, plasma insulin, three-pool model parameters $F_{m,a}$ and $F_{m,0}$, Akt, p70S6K, and MPS following a 3.5-gram bolus of leucine. Data points represent experimental data collected from two studies following the ingestion of a 3.5-gram bolus of leucine in human subjects[1,8]. Root mean square values for each time-course are included within the respective plot. The measured data are presented as means ± SE. Int. = intracellular.

searched the PubMed database to locate studies that measured time-courses of phospho-p70S6K$^{T389}$ in young adults following the ingestion of leucine in non-exercised human subjects. Our search string included the keywords p70S6K, leucine, human, and skeletal muscle but we were unable to find eligible articles. We revisited the studies that were identified during our data acquisition and we were able to locate several articles that met our inclusion criteria. We hand searched the publication history of the primary investigators from the eligible studies to locate additional studies. We extracted the data from each study (e.g., leucine dose, phospho-data fold change, SE) and inputted the data into a spreadsheet. We then meta-analyzed the data using spline regression ('gam' function in R) to fit the extracted data and used the model to predict the time course of phospho-p70S6K at 0, 30, 60, 90, 120, 150, and 180 minutes ('predict.gam' function in R). The same protocol was applied to phospho-Akt$^{S473}$.

Our systematic review of phospho-p70S6K$^{T389}$ time-courses identified six eligible articles that featured seven independent intervention groups of young adults from three distinct research groups [Atherton[2,3], Moore[4,5], Rasmussen[1,6]]. We found that the studies from the Rasmussen lab featured more discrepant fold changes in phospho-p70S6K$^{T389}$ (3.85 – 39.5 fold change, Supplementary Figure C, A) at time 60 minutes in comparison to the data from the other two groups. We were unable to discern from the methods the reasons for these discrepant results, so we removed these

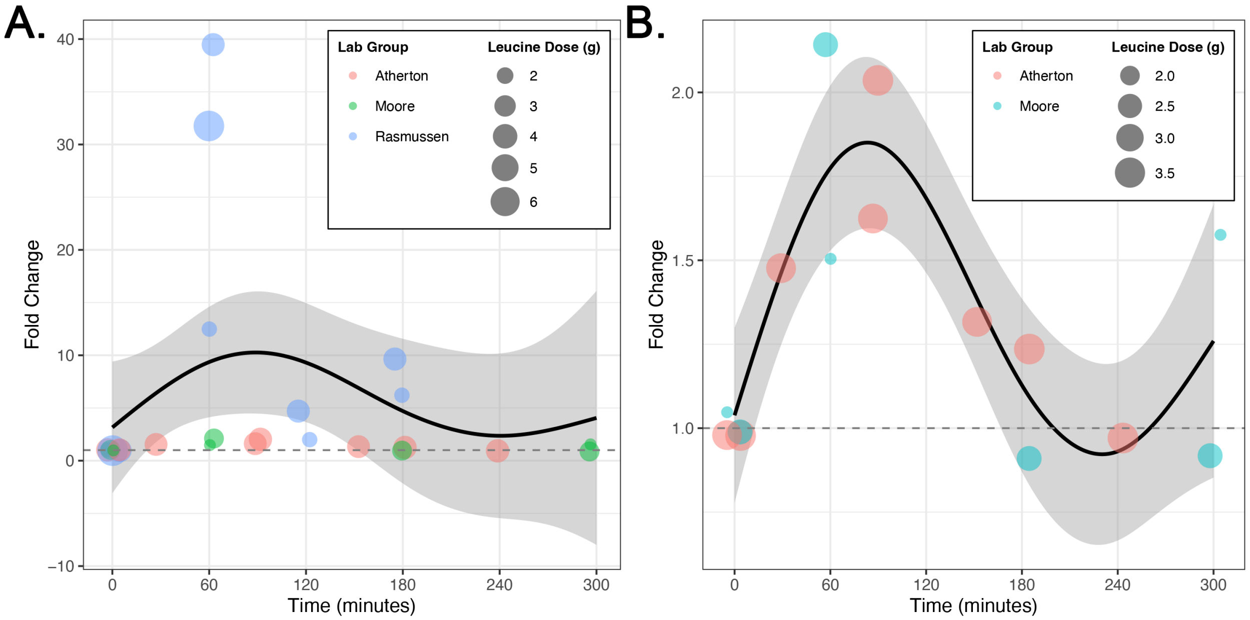


**Figure C. Meta-analysis of phospho-p70S6K$^{T389}$ time course measurements in young adult skeletal muscle following leucine ingestion.**
Scatterplot between phospho-p70S6K$^{T389}$ fold change and time from the data extracted from the scoping review A) with data from the Rasmussen lab group and B) following the removal of data from the Rasmussen lab group. The black line is the predicted phospho-p70S6K$^{T389}$ time-course in response to a 3.5-gram bolus of leucine and the gray areas represents the 95% confidence error of the prediction. The size of the data points corresponds to the leucine dose administered within the intervention. The colour of data points corresponds to the lab group from which the data was reported. Data points are jittered to minimize overlap.

studies from further analysis. Following the removal of these studies, four eligible articles remained that featured four independent intervention groups. Visual inspection of the data from these remaining studies suggested that the fold-changes in phospho-p70S6K$^{T389}$ levels used for calibrating version 1 of the model may have been overestimated[1,6]. We meta-analyzed the data using spline regression to quantify the time course of phospho-p70S6K$^{T389}$ fold changes, with leucine dose included as a covariate (k=4, adjusted $R^2$=0.64, deviance explained=74%, n=15; Supplementary Figure C, panel B). We used the resulting model to predict the time-course of phospho-p70S6K$^{T389}$ in response to a 3.5-gram leucine bolus for future model calibration.

The systematic review of phospho-Akt$^{S473}$ time-courses located five eligible studies that featured five independent intervention groups[1–4,6]. We applied spline regression to quantify the relationship of phospho-Akt$^{S473}$ fold change over time in response to the ingestion of a bolus of leucine, in which leucine dose was included as a covariate (k=5, adjusted $R^2$=0.46, deviance explained=58.1%, n=22; Supplementary Figure D). We used the resulting model to predict the time-course of phospho-Akt$^{S473}$ in response to a 3.5-gram leucine bolus for future model calibration.

*Model development: Version 1 of the model calibrated to the predicted signaling data shows lack-of-fit to validation data*

We repeated model calibration using the predicted phospho-p70S6K$^{T389}$ and phospho-pAkt$^{S473}$ time-courses from the spline regressions in response to ingestion of a 3.5-gram bolus of leucine. Calibration of the model parameters resulted in a root mean square value of 6.71. The model simulations qualitatively agreed with experimental data for plasma leucine, intracellular leucine, $F_{m,a}$, $F_{m,0}$, phospho-Akt$^{S473}$, phospho-p70S6K$^{T389}$, and MPS in response to a 3.5-gram bolus of leucine (Supplementary Figure E). Plasma insulin exhibited a reasonable dynamic (i.e., timing of peak concentration) but had a reduced peak concentration compared to the experimental data.

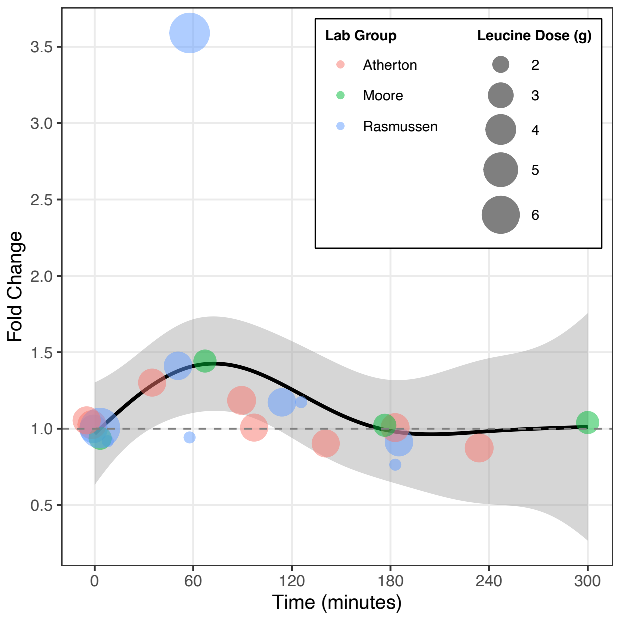


**Figure D. Meta-analysis of phospho-Akt$^{S473}$ time course measurements in young adult skeletal muscle following leucine ingestion.** Scatterplot between phospho-Akt$^{S473}$ fold change and time from the data extracted from the scoping review. The black line is the predicted phospho- Akt$^{S473}$ time-course in response to a 3.5-gram bolus of leucine and the gray areas represents the 95% confidence error of the prediction. The size of the data points corresponds to the leucine dose administered within the intervention. The colour of data points corresponds to the lab group from which the data was reported. Data points are jittered to minimize overlap.

Next, we aimed to validate version 1 of the model by comparing the output of the calibrated model against data from a pulsatile feeding protocols [2] and five bolus feeding protocols[1–3,7,8]. The pulsatile feeding protocol was simulated by providing four 0.9-gram doses of leucine at 45-minute intervals, whereas the bolus feeding protocols were simulated by providing a 1.85-gram, 3,42-gram, 3.5-gram, or 3.59-gram bolus of leucine at 0 minutes. The pulsatile feeding protocol was markedly different from the experimental data used to calibrate the model, such that it served as a stringent test for model validation. In simulating the pulsatile and bolus feeding protocols, only the dose and timing of the leucine dose was varied. The model prediction for the validation data sets generally qualitatively agreed with plasma leucine, intracellular leucine, phospho-Akt$^{S473}$, phospho-p70S6K$^{T389}$, and MPS. However, the model underestimated plasma insulin, in particular the early phase of the dynamic where the peak insulin concentration occurred (~30 minutes) (Supplementary Figure F). Additionally, the model predicted cumulative increases in plasma

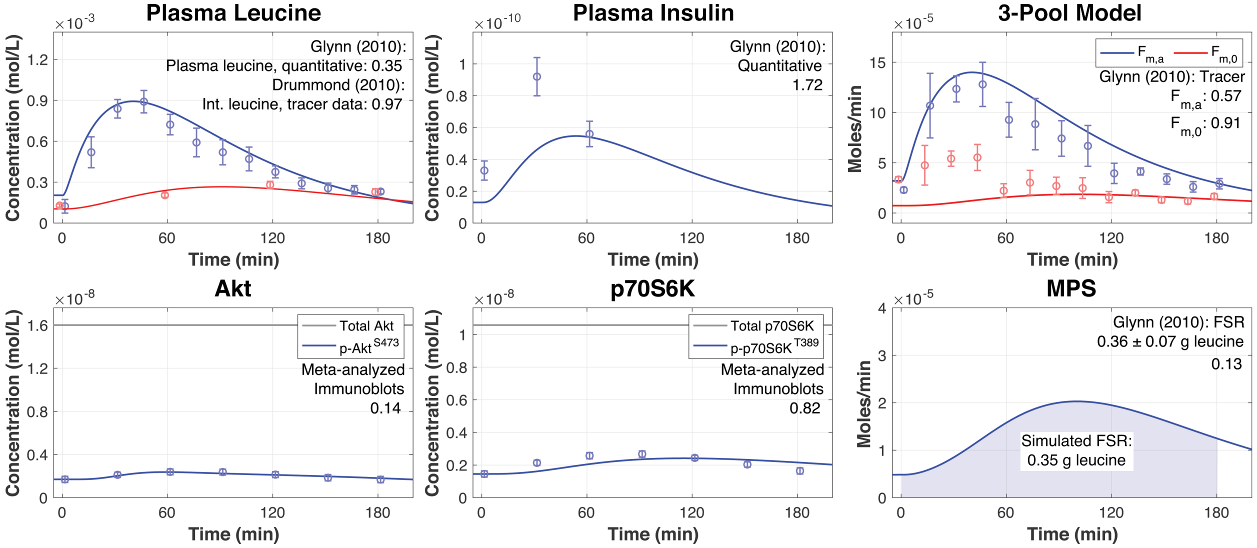


**Figure E. Model calibration of model version 1 using the meta-analyzed phospho-data.** Simulated time-courses following model version 1 calibration of plasma leucine, intracellular leucine, plasma insulin, three-pool model parameters $F_{m,a}$ and $F_{m,0}$, Akt, p70S6K, and MPS following a 3.5-gram bolus of leucine. Data points represent experimental data collected from two studies following the ingestion of a 3.5-gram bolus of leucine in human subjects[1,8]. The data points for phospho-Akt$^{S473}$ and phopsho-p70S6K$^{T389}$ were predicted from the meta-analyzed spline regressions. Root mean square values for each time-course are included within the respective plot. The measured data are presented as means ± SEM. Int. = intracellular.

insulin concentration following pulsatile leucine feeding, whereas the experimental data showed a sustained plasma insulin concentration.

We posited that this lack of fit reflected the influence of a negative feedback mechanism that blunted further insulin release in response to subsequent feedings, which was not represented in the current model.

*Model development: Including an insulin secretion module to simulate the oscillatory secretion of insulin*

We searched the literature to identify feedback mechanisms that could produce physiological insulin dynamics and located a model that simulates insulin secretion[9]. Following leucine ingestion, insulin is secreted from the pancreas [10,11] and this occurs in an oscillatory pattern with a period of 80-150 minutes (i.e., ultradian), with more rapid pulses of insulin secretion with a period of 10-15 minutes that are overlaid on the ultradian oscillatory insulin secretion[9]. The Tolic et al.[9] model simulates the ultradian oscillations of insulin secretion using two feedback loops between glucose and insulin: 1) the effect of insulin on glucose utilization and 2) the effect of insulin on glucose production. Both feedback loops also include the stimulatory effect of glucose on insulin secretion. The Tolic et al.[9] model includes three compartments (plasma insulin, intracellular insulin, plasma glucose) and three variables that represent the delay between plasma insulin and its effect on hepatic glucose production.

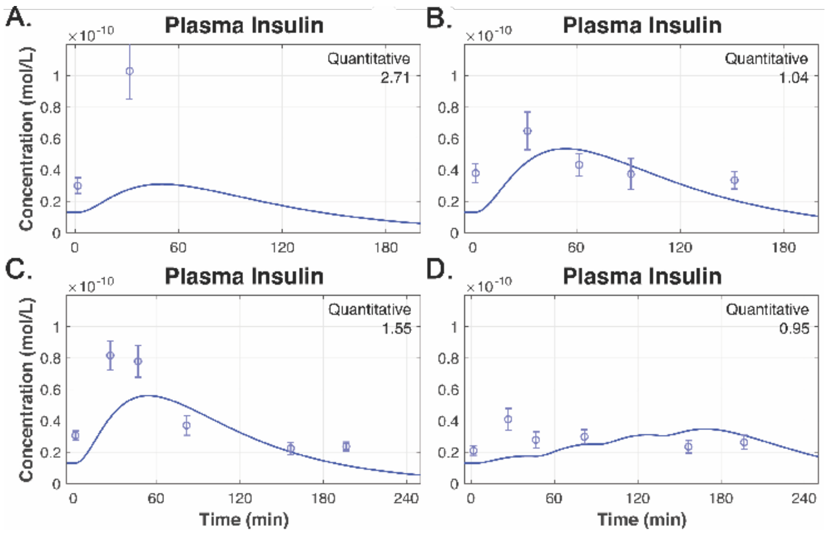


**Figure F. Version 1 of the model inaccurately predicts the early dynamics of insulin in response to leucine feedings.**
A-C) Simulated plasma insulin time-courses in response to different bolus leucine feeding (1.85-grams, 3.42-grams, and 3.5-grams, respectively). Data points in A), B), and C) represent experimental data from Glynn et al.[1], Wilkinson et al.[3], and Mitchell et al.[2], respectively. D) Simulated plasma insulin time-course in response to pulsatile leucine feeding (i.e., four 0.9-gram boluses of leucine administered at times 0, 45, 90, and 135 minutes). Data points represent experimental data from Mitchell et al.[2]. Root mean square values for each time-course are included within the respective plot. The measured data are presented as means ± SEM.

We included the Tolic et al.[9] module in our model in place of the insulin variable and added a leucine mediated insulin secretion kinetic reaction (reaction 4; model version 2). This reaction increases the plasma insulin content dependent on the concentration of plasma leucine. Our simulation does not directly affect plasma glucose concentration, but changes in plasma insulin affect plasma glucose. The interplay between plasma insulin and plasma glucose leads to the oscillatory dynamic observed in the model. Inclusion of the Tolic et al.[9] module allowed for a more accurate simulation of basal insulin values at time 0 and better simulated the timing of peak insulin concentrations (Figure 2, Supplementary Figure G).

Model version 2 includes the mTOR signalling module from Dalle Pezze et al.[12], the leucine kinetic module from Tessari et al.[9], the digestive system module that we developed, and the insulin secretion module from Sturis et al.[13]. Model version 2 is henceforth referred to as 'the model' in the main text.

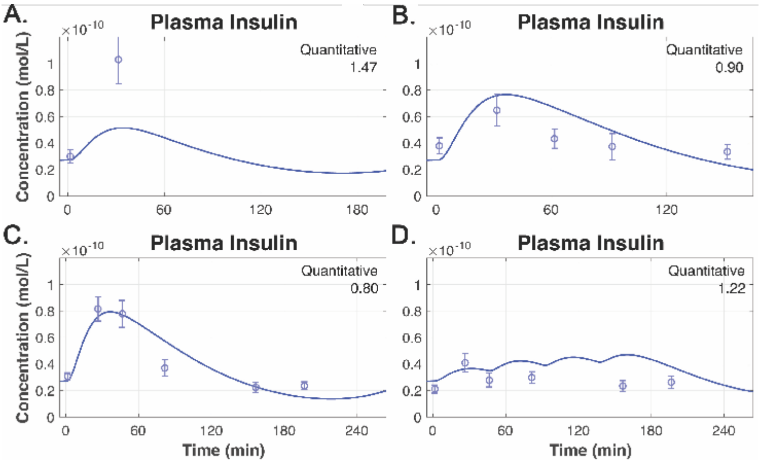


**Figure G. Version 2 of the model better simulates the plasma insulin in response to leucine feedings.**

A-C) Simulated plasma insulin time-courses in response to different bolus leucine feeding (1.85-grams, 3.42-grams, and 3.5-grams, respectively). Data points in A), B), and C) represent experimental data from Glynn et al. (2010) (25), Wilkinson et al. (2013) (52), and Mitchell et al. (2015) (33), respectively. D) Simulated plasma insulin time-course in response to pulsatile leucine feeding (i.e., four 0.9-gram boluses of leucine administered at times 0, 45, 90, and 135 minutes). Data points represent experimental data from Mitchell et al. (2015) (33). Root mean square values for each time-course are included within the respective plot. The measured data are presented as means ± SE.

10. Newsholme, P., Brennan, L., and Bender, K. (2006). Amino Acid Metabolism, β-Cell Function, and Diabetes. Diabetes *55*, S39–S47. 10.2337/db06-S006.

11. Newsholme, P., Cruzat, V., Arfuso, F., and Keane, K. (2014). Nutrient regulation of insulin secretion and action. The Journal of endocrinology *221*, R105-20. 10.1530/JOE-13-0616.

12. Dalle Pezze, P., Sonntag, A.G., Thien, A., Prentzell, M.T., Gödel, M., Fischer, S., Neumann-Haefelin, E., Huber, T.B., Baumeister, R., Shanley, D.P., et al. (2012). A dynamic network model of mTOR signaling reveals TSC-independent mTORC2 regulation. Science Signaling *5*. 10.1126/scisignal.2002469.

13. Sturis, J., Polonsky, K.S., Mosekilde, E., and Van Cauter, E. (1991). Computer model for mechanisms underlying ultradian oscillations of insulin and glucose. The American journal of physiology *260*, E801-9. 10.1152/ajpendo.1991.260.5.E801.
