## Supplementary figures and tables for "Kinetic modeling of leucine-mediated signaling and protein metabolism in human skeletal muscle"

### Contents:


Figure S1. Related to Figure 2. Comprehensive model validation.

Figure S2. Related to Figure 2. Discrepant plasma leucine measurements between different data sets

Figure S3. Related to Figure 2. Simulation of all unmeasured species in the model

Figure S4. Related to Figure 3. Model simulation following the oral ingestion of rapamycin, a potent mTORC1 inhibitor.

Figure S5. Related to Figure 3. Phospho-p70S6K signaling is required to elicit a maximal MPS response.

Table S1. Related to Figure 2. Proteins included in the model and their cellular properties.

Table S2. Related to Figure 2. Reaction rate equations.

Table S3. Related to Figure 2. System of ordinary differential equations.

Table S4. Related to Figure 2. Model species and initial concentrations.

Table S5. Related to Figure 2. Model parameter values.

Table S6. Related to Figure 2A. Calibration dataset characteristics.

Table S7. Related to Figure 2B,C. Validation dataset characteristics.

Table S8. Related to Figure 2. Amino acid profile for each intervention.

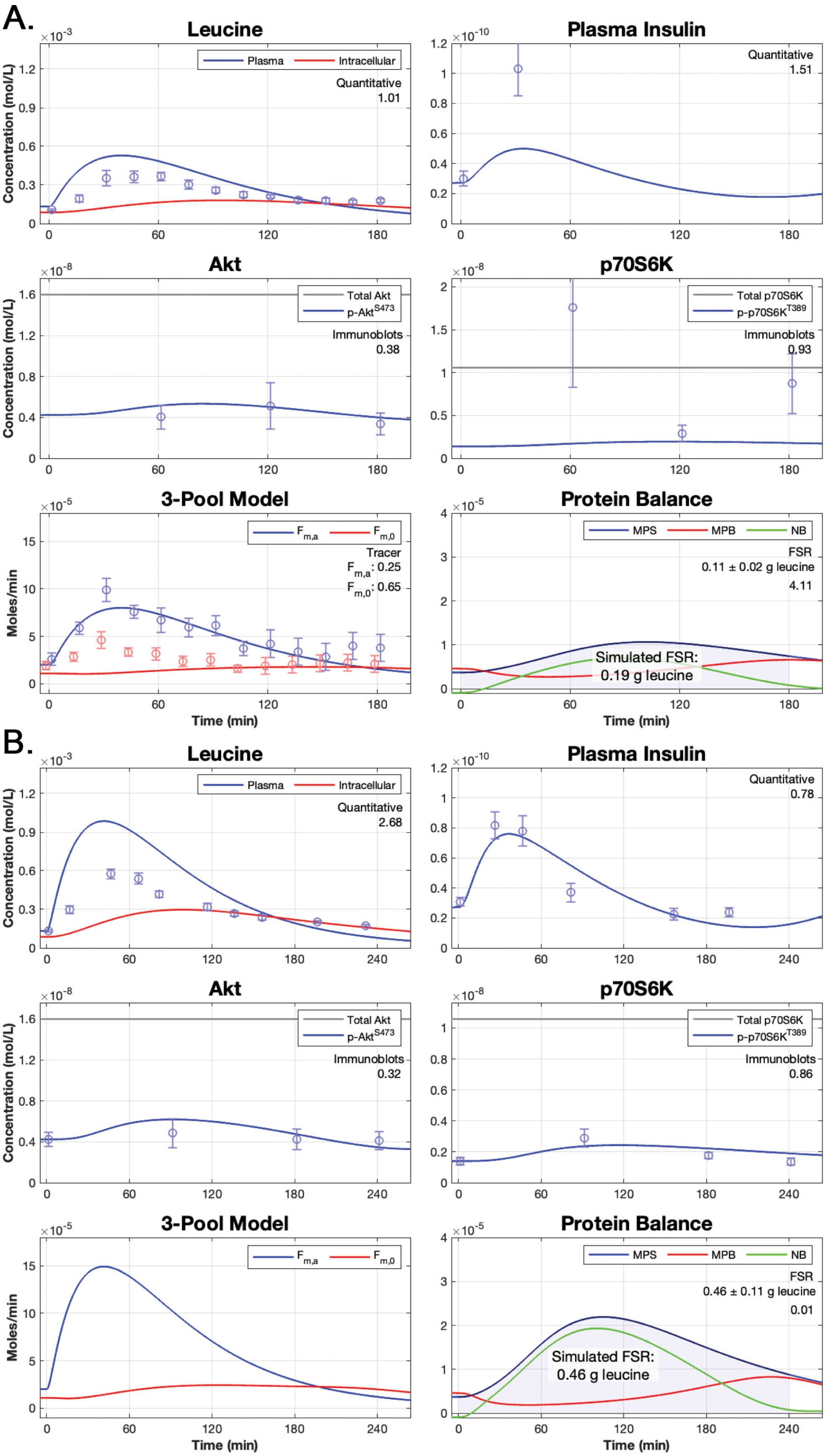
A.
Leucine
Plasma
Intracellular
Quantitative
1.01
Concentration (mol/L)
Plasma Insulin
Quantitative
1.51
Akt
Total Akt
p-Akt$^{S473}$
Immunoblots
0.38
p70S6K
Total p70S6K
p-p70S6K$^{T389}$
Immunoblots
0.93
3-Pool Model
$F_{m,a}$
$F_{m,0}$
Tracer
$F_{m,a}$: 0.25
$F_{m,0}$: 0.65
Moles/min
Time (min)
Protein Balance
MPS
MPB
NB
FSR
0.11 ± 0.02 g leucine
4.11
Simulated FSR:
0.19 g leucine
B.
Leucine
Plasma
Intracellular
Quantitative
2.68
Plasma Insulin
Quantitative
0.78
Akt
Total Akt
p-Akt$^{S473}$
Immunoblots
0.32
p70S6K
Total p70S6K
p-p70S6K$^{T389}$
Immunoblots
0.86
3-Pool Model
$F_{m,a}$
$F_{m,0}$
Protein Balance
MPS
MPB
NB
FSR
0.46 ± 0.11 g leucine
0.01
Simulated FSR:
0.46 g leucine

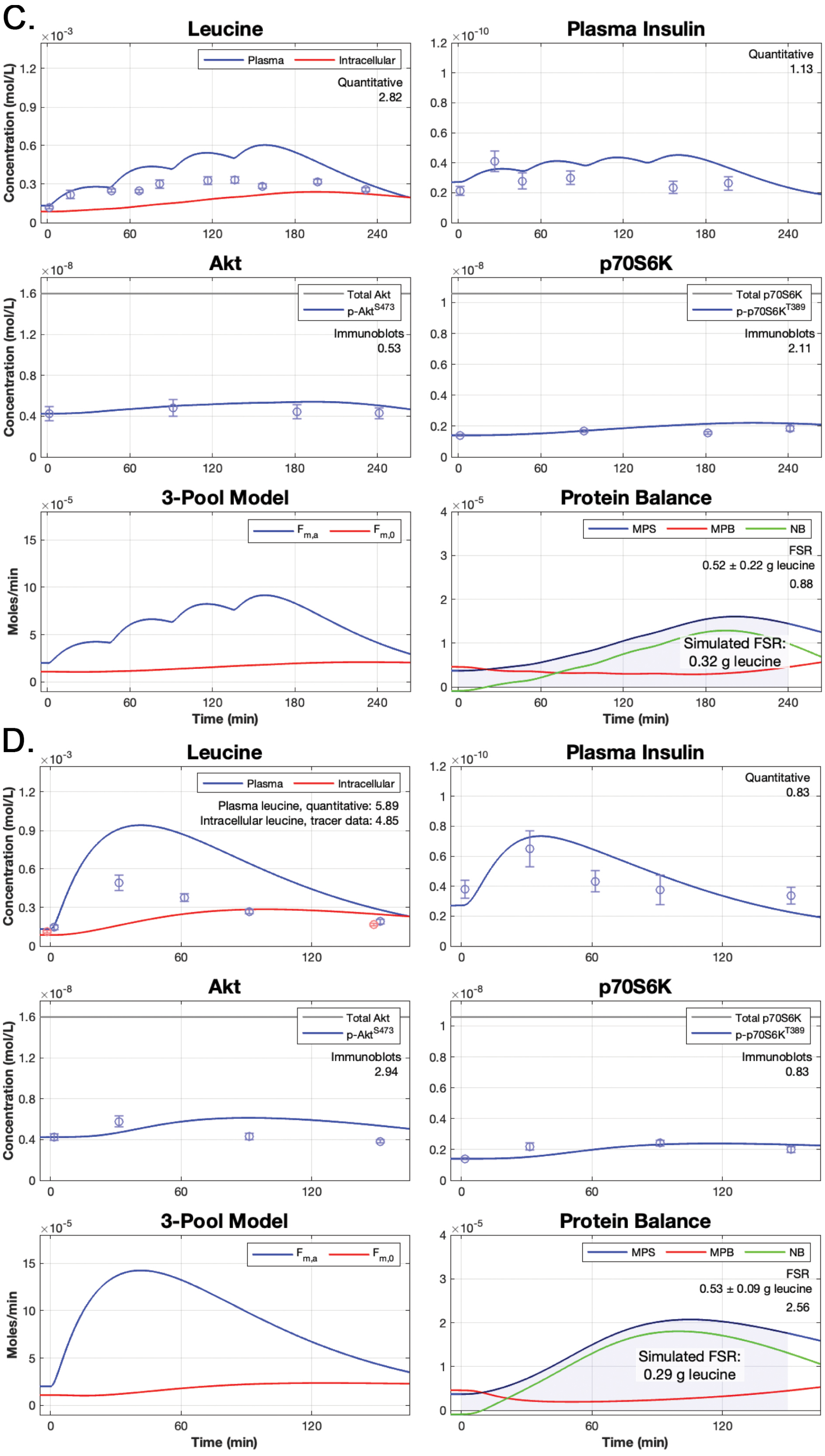
C.
Leucine
Plasma
Intracellular
Quantitative
2.82
Concentration (mol/L)
Plasma Insulin
Quantitative
1.13
Akt
Total Akt
p-Akt S473
Immunoblots
0.53
p70S6K
Total p70S6K
p-p70S6K T389
Immunoblots
2.11
3-Pool Model
Moles/min
F m,a
F m,0
Time (min)
Protein Balance
MPS
MPB
NB
FSR
0.52 ± 0.22 g leucine
0.88
Simulated FSR:
0.32 g leucine
D.
Leucine
Plasma
Intracellular
Plasma leucine, quantitative: 5.89
Intracellular leucine, tracer data: 4.85
Plasma Insulin
Quantitative
0.83
Akt
Total Akt
p-Akt S473
Immunoblots
2.94
p70S6K
Total p70S6K
p-p70S6K T389
Immunoblots
0.83
3-Pool Model
F m,a
F m,0
Protein Balance
MPS
MPB
NB
FSR
0.53 ± 0.09 g leucine
2.56
Simulated FSR:
0.29 g leucine

E.

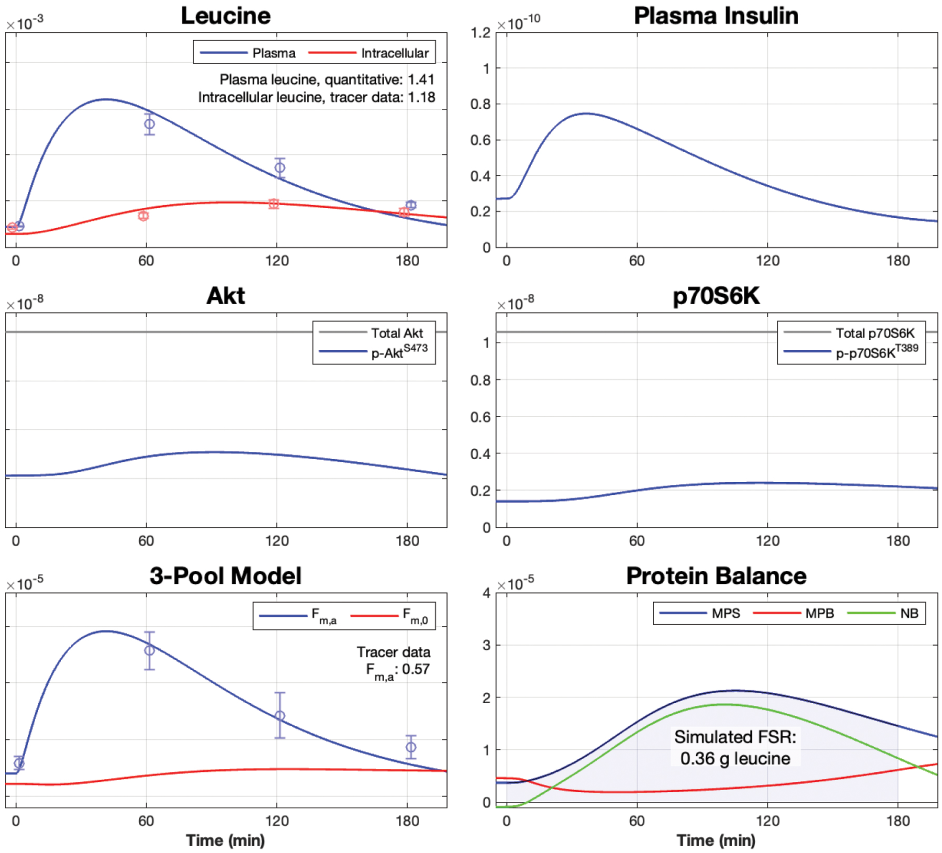
Leucine
Plasma
Intracellular
Plasma leucine, quantitative: 1.41
Intracellular leucine, tracer data: 1.18
Concentration (mol/L)
Plasma Insulin
Akt
Total Akt
p-Akt$^{S473}$
p70S6K
Total p70S6K
p-p70S6K$^{T389}$
3-Pool Model
$F_{m,a}$
$F_{m,0}$
Tracer data
$F_{m,a}$: 0.57
Moles/min
Time (min)
Protein Balance
MPS
MPB
NB
Simulated FSR:
0.36 g leucine
Time (min)

F.

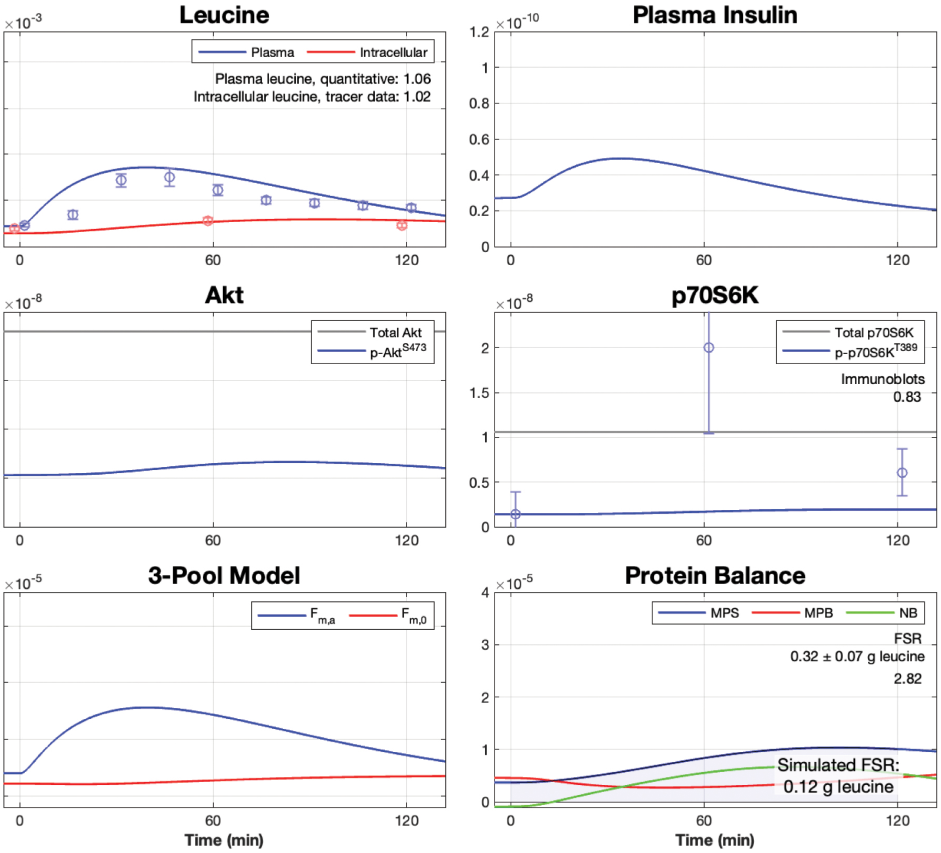
Leucine
Plasma
Intracellular
Plasma leucine, quantitative: 1.06
Intracellular leucine, tracer data: 1.02
Concentration (mol/L)
Plasma Insulin
Akt
Total Akt
p-Akt$^{S473}$
p70S6K
Total p70S6K
p-p70S6K$^{T389}$
Immunoblots
0.83
3-Pool Model
$F_{m,a}$
$F_{m,0}$
Moles/min
Time (min)
Protein Balance
MPS
MPB
NB
FSR
0.32 ± 0.07 g leucine
2.82
Simulated FSR:
0.12 g leucine
Time (min)

**Figure S1. Related to Figure 2. Comprehensive model validation.**
Simulated time-courses of plasma leucine, plasma insulin, Akt, p70S6K, three-pool model parameters $F_{m,a}$ and $F_{m,0}$, and muscle protein balance following a single (A) 1.85-gram, (B) 3.59-gram, (D) 3.42-gram, (E) 3.50-gram, or (F) 1.80-gram bolus of leucine or (C) pulsatile leucine feedings (0.59-grams of leucine provided at 0, 45, 90, and 135 minutes). Data points represent experimental data collected from (A) Glynn et al.[1], (B,C) Mitchell et al.[2], (D) Wilkinson et al.[3], (E) Drummond et al.[4], and (F) Dickinson et al[5]. Root mean square values for each time-course with corresponding experimental data are included within the respective plot. The measured data are presented as means ± SE.

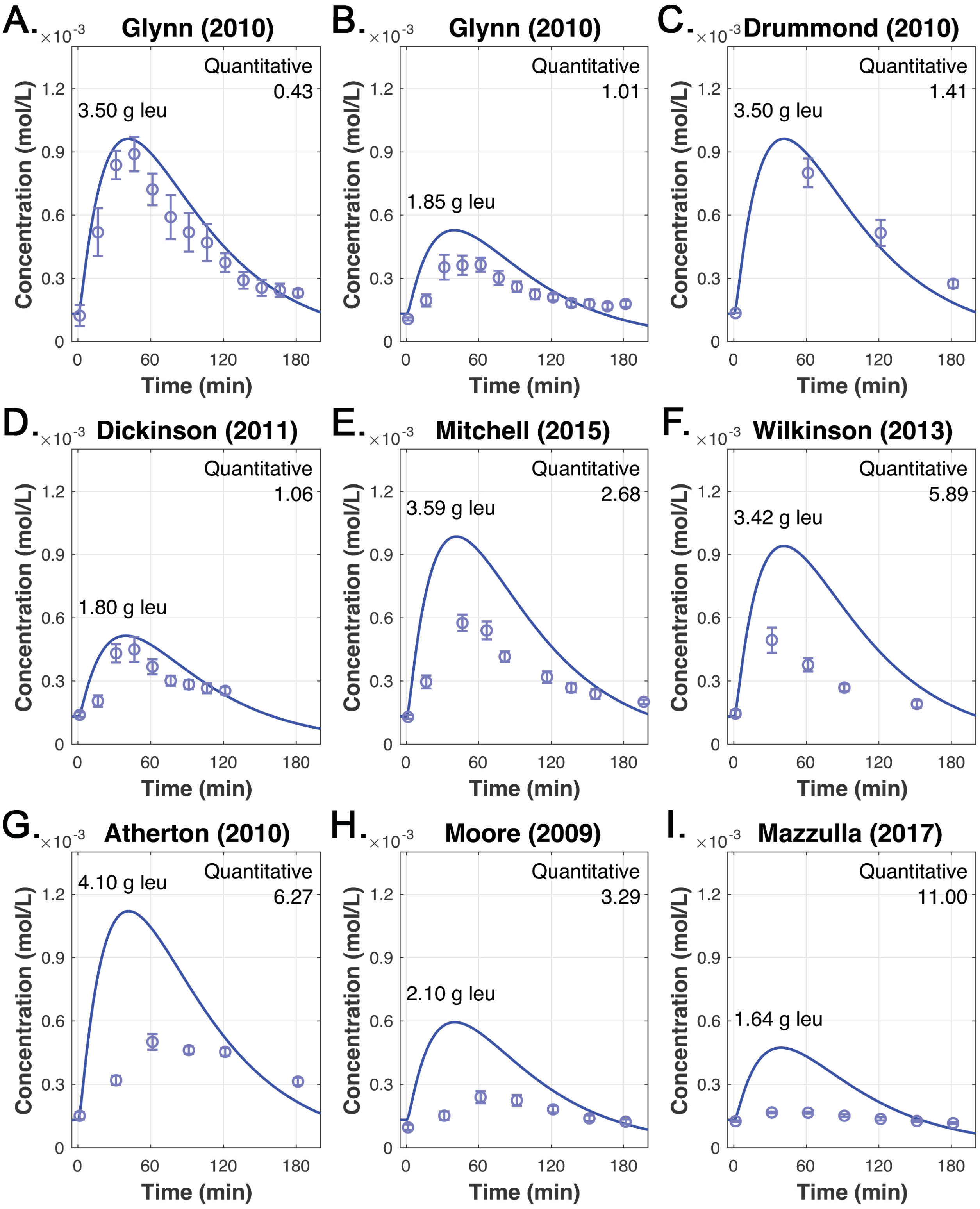


**Figure S2. Related to Figure 2. Discrepant plasma leucine measurements between different data sets**

Simulated plasma leucine time-courses in response to bolus leucine feedings (A-F: 3.5 g, 1.85 g, 3.50 g, 1.80 g, 3.59 g, and 3.42 g, respectively) and whey protein feedings with differing leucine amounts (G-I: 4.1 g, 2.1 g, or 1.67 g, respectively). Data points represent experimental data from

(A) Glynn et al.[1], (B) Glynn et al.[1], (C) Drummond et al.[4], (D) Dickinson et al.[5], (E) Mitchell et al.[2], (F) Wilkinson et al.[3], (G) Atherton et al.[6], (H) Moore et al.[7], and (I) Mazzulla et al.[8]. The ingested dose of leucine is listed on the left side of each plot and the root mean square values for each time-course is listed in the upper-right portion of each plot. The measured data are presented as means ± SE.

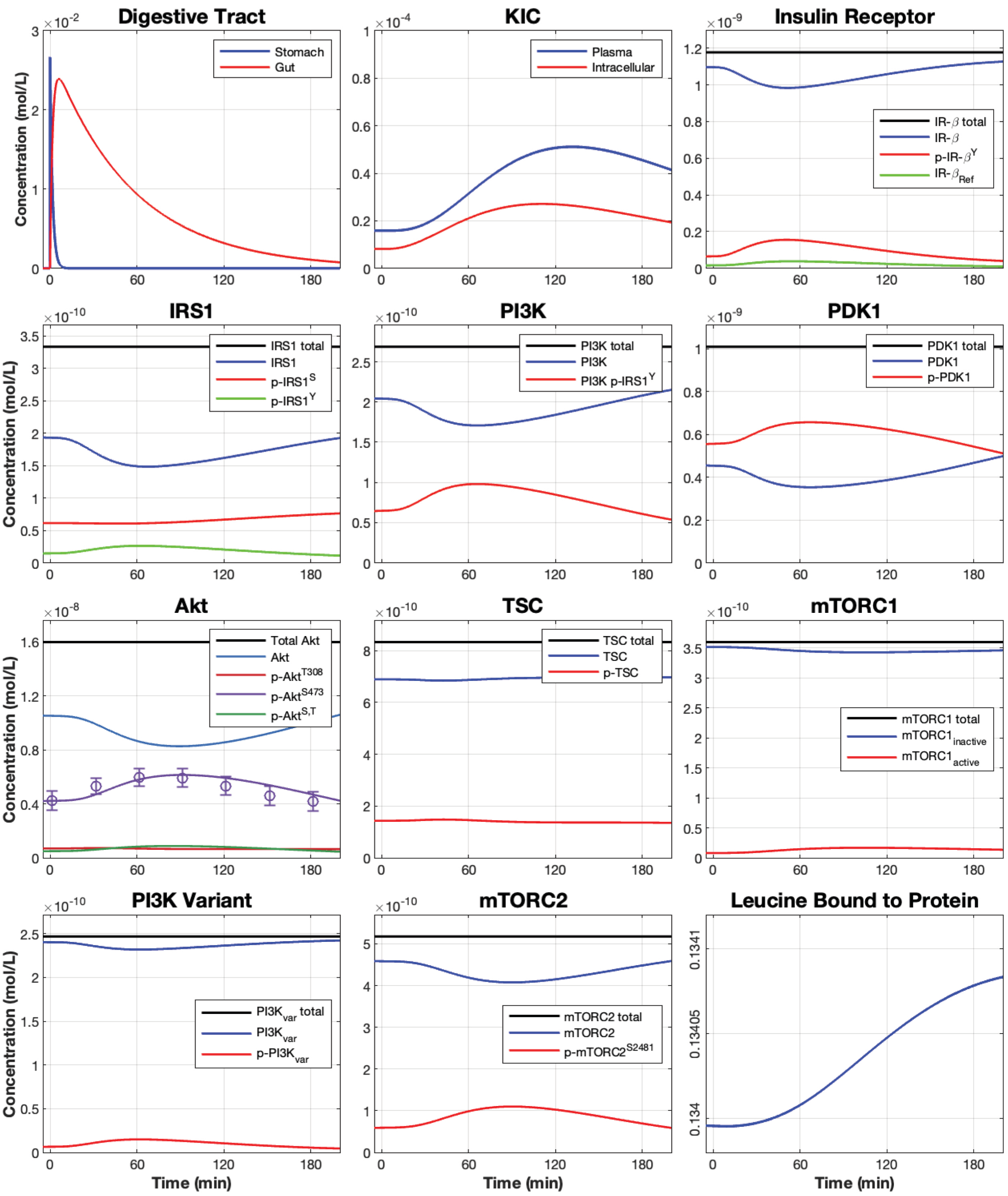


**Figure S3. Related to Figure 2. Simulation of unmeasured species in the model**

Simulated time-courses of all model species that were not experimentally measured following a 3.5-gram bolus of leucine. The phospho-Akt$^{S473}$ time-course includes the predicted time-course data from the meta-analyzed spline regression (mean ± SE). Total IRS1 includes the PI3K p-IRS1$^{Y}$ complex, which is presented in the adjacent PI3K panel.

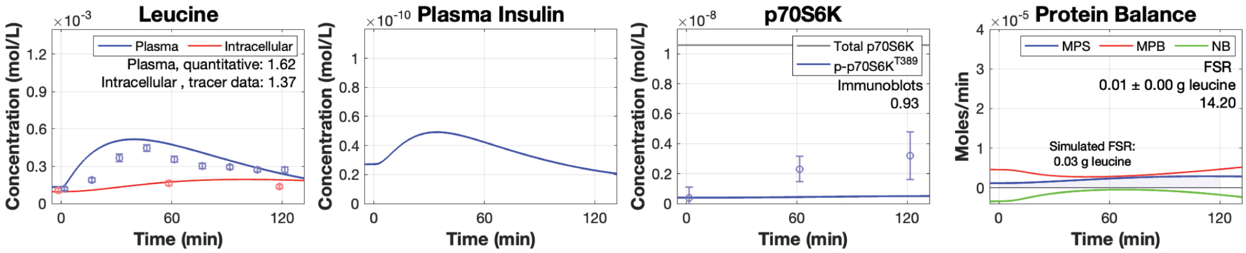


**Figure S4. Related to Figure 3. Model simulation of rapamycin ingestion.**

Simulated time-courses of plasma leucine, intracellular leucine, plasma insulin, p70S6K, and muscle protein balance following a 1.8-gram bolus of leucine with prior ingestion of rapamycin, a potent mTORC1 inhibitor. Data points represent experimental data from Dickinson et al.[5]. Root mean square values for each time-course with corresponding experimental data are included within each plot. The measured data are presented as means ± SE.

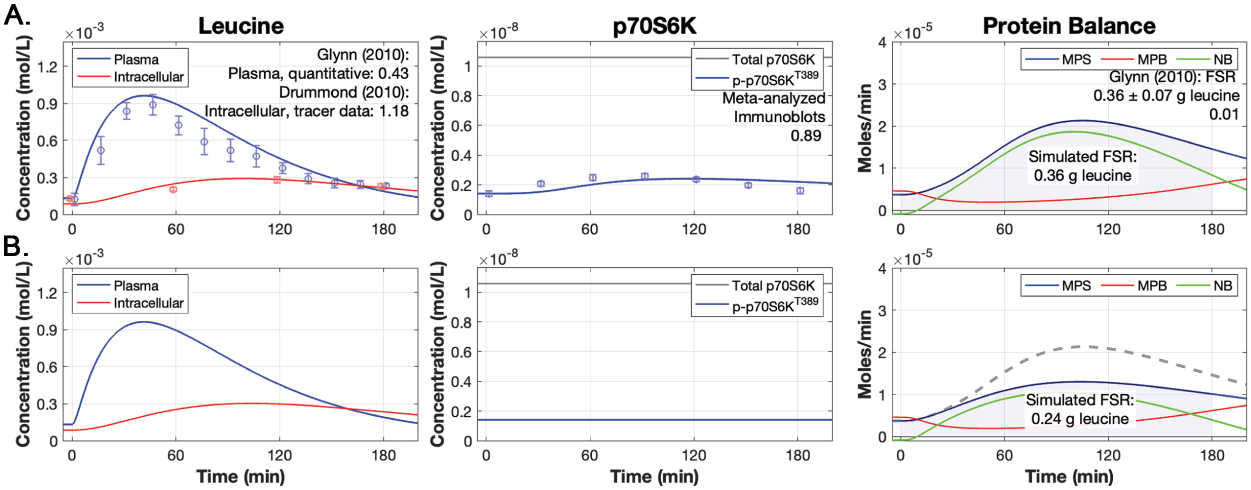


**Figure S5. Related to Figure 3. Phospho-p70S6K signaling is required to elicit a maximal MPS response.**

Simulated plasma leucine, intracellular leucine, p70S6K, and muscle protein balance time courses following a 3.5-gram bolus of leucine with (A) phospho-p70S6K sensitive to leucine-mediated signaling or (B) phospho-p70S6K insensitive to leucine-mediated signaling and maintained at the calibrated concentration over the simulation duration. Data points in (A) represent data from the model calibration. The gray dashed line in the (B) protein balance plot is a reference line for the original MPS from (A). Root mean square values for each time-course with corresponding experimental data are included within each plot. The measured data are presented as means ± SE.

**Table S1. Related to Figure 2. Proteins included in the model and their cellular properties.**

| Species | Name | Properties | Ref. |
|---|---|---|---|
| Akt | Protein kinase B | Akt is an AGC kinase that must be phosphorylated twice to achieve full activity. Akt translocates to the plasma membrane by interacting with $PIP_3$, where PDK1 phosphorylates the Thr308 residue of Akt and mTORC2 phosphorylates the Ser473 residue of Akt. Thr308 and Ser473 may be independently phosphorylated such that there are two pathways that can achieve full activation of Akt. $Akt^{T308}$ and $Akt^{T308,S473}$ function to phosphorylate the TSC1/2 complex. | [9–13] |
| $IR_{\beta}$ | Insulin receptor beta | Insulin binds to the IR, which leads to the phosphorylation of the Tyr1164 site and other tyrosine residues. Tyrosine phosphorylated IR promotes phosphorylation of the IRS1 tyrosine residues. | [10,14] |
| IRS1 | Insulin receptor substrate 1 | IRS1 can be phosphorylated at two sites: Ser636 and Tyr residue. The Tyr residue of IRS1 is phosphorylated by $IR_{\beta}^{Y1164}$ and functions to activate the PI3K species. The Ser636 species is phosphorylated by $p70S6K^{T389}$, which disrupts the ability of IRS1 to interact with activated $IR_{\beta}$. Phosphorylation of the Ser636 residue denotes a negative feedback loop mediated by p70S6K that inhibits insulin-mediated mTORC1 activity. | [10,14–18] |
| mTORC1 | Mechanistic target of rapamycin complex 1 | mTORC1 activity is mediated through insulin- and leucine-independent pathways. Insulin activates mTORC1 through a signalling cascade that promotes TSC1/2 phosphorylation to inactivate TSC1/2 inhibition. Removal of TSC inhibition allows mTORC1 to translocate to the lysosome where it is activated by Rheb. Leucine activates the Ragulator-Rag complex that directs mTORC1 to the surface of lysosome where it is activated by Rheb. mTORC1 mediates protein translational dynamics by phosphorylating downstream proteins (e.g., p70S6K1, 4EBP1). mTORC1 is inhibited by PRAS40, which interacts with RAPTOR to negatively regulate kinase activity. Additionally, p70S6K regulates mTORC1 activity via a negative feedback loop in which it phosphorylates the Ser2448 residue. | [5,10,19–25] |

| | | | |
|---|---|---|---|
| mTORC2 | Mechanistic target of rapamycin complex 2 | mTORC2 resides at the plasma membrane, is regulated by growth factors, and phosphorylates the Ser473 residue of Akt. In response to growth factors, mTORC2 is phosphorylated at the Ser2481 residue, which promotes phosphorylation of the Akt Ser473 residue. | 10,17,26 |
| p70S6K | p70 ribosomal protein S6 kinase | p70S6K1 is a serine/threonine kinase whose maximum activation requires phosphorylation of the Thr389 and Thr229 residues by mTORC1 and PDK1, respectively. p70S6K1 promotes translational control primarily by phosphorylating the proteins eukaryotic initiation factor (eIF) 4B, eIF3, and PDCD4 (programmed cell death protein 4). Additionally, p70S6K1 controls protein elongation by phosphorylating eukaryotic elongation factor (eEF) 2K and eEF1A [27]. Like p70S6K1, 4EBP1 controls translation by controlling initiation, ribosomal biogenesis, and export of mRNA from the nucleus. p70S6K participates in several feedback loops that function to reduce mTORC1 activity (e.g., p70S6K/IRS1, p70S6K/mTORC1). | 5,10,16,20, 22,23,27–30 |
| PDK1 | 3-phosphoinositide dependent kinase-1 | PDK1 is activated by PI3K and functions to phosphorylate the Thr308 residue of Akt and the Thr229 residue of p70S6K. | 9,10,28,30, 31 |
| PI3K | Phosphoinositide 3-kinase | Tyrosine phosphorylated IRS1 serves as a docking site for the p85 regulatory subunit of PI3K, which results in the activation of PI3K. Activated PI3K phosphorylates PI(4,5)P2 to form PIP3 that activates PDK1. | 9,10,14,32, 33 |
| $PI3K_{variant}$ | PI3K variant | A wortmannin-sensitive, but IRS1 independent PI3K species proposed by Dalle Pezze et al. that is activated by $IR_{\beta}$ and stimulates mTORC2 in response to insulin. | 12 |
| TSC1/2 | Tuberous sclerosis protein 1/2 | The TSC1/2 complex negatively regulates mTORC1 activity. The complex is phosphorylated by $Akt^{T308}$ and $Akt^{T308,S473}$, which inactivates the inhibitory function of the complex. This inactivation of the inhibitory function promotes the RHEB GTPase to form the active RHEB/GTP complex, which activates mTORC1. | 10,12,34,35 |

**Table S2. Related to Figure 2. Reaction rate equations.**
Reaction rate equations and the functional relationships used in the oscillatory insulin module.

| **Description of reaction** | **Reaction** | **Reaction rates** |
|---|---|---|
| | *Digestive module* | |
| Leucine transport from stomach to gut | Stomach → Gut | $r1 = k1 \times Leucine_{stomach}$ |
| Leucine excretion via the gut | Gut → ∅ | $r2 = k2 \times Leucine_{gut}$ |
| First pass splanchnic extraction | Gut → ∅ | $r68 = k68 \times Leucine_{gut}$ |
| Absorption of leucine from the gut to the blood plasma | Gut → Plasma leucine | $r3 = k3 \times Leucine_{gut}$ |
| | *Blood Plasma* | |
| Leucine mediated insulin secretion | Plasma leucine → Plasma insulin | $r4 = k4 \times Leucine_{plasma}$ |
| | *Leucine kinetic module* | |
| Leucine transport into skeletal muscle cells | Plasma leucine → Intracellular leucine | $r6 = k6 \times Leucine_{plasma}$ |
| Leucine release from skeletal muscle to blood plasma | Intracellular leucine → Plasma leucine | $r7 = k7 \times (Leucine_{intracellular} - Leucine_{plasma})$ |
| Protein synthesis | Intracellular leucine → Protein | $r15 = k15 \times Leucine_{intracellular} \times p70S6K^{T389}$ |
| Protein breakdown | Protein → Intracellular leucine | $r9 = k9 \times Protein \times \frac{1}{p\text{-}IR\text{-}\beta^{Y1164}}$ |
| Leucine transamination to KIC | Intracellular leucine → Intracellular KIC | $r10 = k10 \times Leucine_{intracellular}$ |

| | | |
|---|---|---|
| KIC reamination to leucine | Intracellular KIC → Intracellular leucine | $r11 = k11 \times KIC_{intracellular}$ |
| KIC transport into skeletal muscle cells | Plasma KIC → Intracellular KIC | $r12 = k12 \times KIC_{Plasma}$ |
| KIC release from skeletal muscle to blood plasma | Intracellular KIC → Plasma KIC | $r13 = k13 \times KIC_{intracellular}$ |
| KIC oxidation | Intracellular KIC → ∅ | $r14 = k14 \times KIC_{intracellular}$ |
| *mTOR signalling module* | | |
| Insulin-mediated IR-$\beta$ phosphorylation | IR-$\beta$ → p-IR-$\beta^{Y1164}$ | $r16 = k16 \times IR\text{-}\beta \times Insulin_{plasma}$ |
| Dephosphorylation of p-IR-$\beta^{Y1164}$, refractory state | p-IR-$\beta^{Y1164}$ → IR-$\beta_{refractory}$ | $r17 = k17 \times p\text{-}IR\text{-}\beta^{Y1164}$ |
| Dephosphorylation of p-IR-$\beta^{Y1164}$, non-refractory state | IR-$\beta_{refractory}$ → IR-$\beta$ | $r18 = k18 \times IR\text{-}\beta_{refractory}$ |
| Phosphorylation of the IRS1 tyrosine site | IRS1 → p-IRS1$^{Y}$ | $r19 = k19 \times IRS1 \times p\text{-}IR\text{-}\beta^{Y1164}$ |
| Dephosphorylation of p-IRS1$^{Y}$ | p-IRS1$^{Y}$ → IRS1 | $r20 = k20 \times p\text{-}IRS1^{Y}$ |
| Phosphorylation of the IRS1 serine site | IRS1 → p-IRS1$^{S}$ | $r21 = k21 \times IRS1 \times p\text{-}p70S6K1^{T389}$ |
| Dephosphorylation of the p-IRS1$^{S}$ | p-IRS1$^{S}$ → IRS1 | $r22 = k22 \times p\text{-}IRS1^{S}$ |
| Docking of PI3K with p-IRS1$^{Y}$ | p-IRS1$^{Y}$ + PI3K → p-IRS1$^{Y}$/PI3K$_{clx}$ | $r23 = k23 \times p\text{-}IRS1^{Y} \times PI3K$ |
| Removal of PI3K from p-IRS1$^{Y}$ | p-IRS1$^{Y}$/PI3K$_{clx}$ → p-IRS1$^{Y}$ + PI3K | $r24 = k24 \times p\text{-}IRS1^{Y}\_PI3K_{clx}$ |
| Phosphorylation of PDK1 | PDK1 → p-PDK1 | $r25 = k25 \times PDK1 \times p\text{-}IRS1^{Y}\_PI3K_{clx}$ |

| | | |
|---|---|---|
| Dephosphorylation of p-PDK1 | p-PDK1 → PDK1 | $r26 = k26 \times p\text{-}PDK1$ |
| Phosphorylation of Akt at the T308 residue | Akt → p-$Akt^{T308}$ | $r27 = k27 \times Akt \times p\text{-}PDK1$ |
| Dephosphorylation of p-$Akt^{T308}$ | p-$Akt^{T308}$ → Akt | $r28 = k28 \times p\text{-}Akt^{T308}$ |
| Phosphorylation of Akt at the S473 residue | Akt → p-$Akt^{S473}$ | $r29 = k29 \times Akt \times p\text{-}mTORC2^{S2481}$ |
| Dephosphorylation of p-$Akt^{S473}$ | p-$Akt^{S473}$ → Akt | $r30 = k30 \times p\text{-}Akt^{S473}$ |
| Phosphorylation of Akt at the S473 residue to form the dual-phosphorylated species | p-$Akt^{T308}$ → p-$Akt^{T308,S473}$ | $r31 = k31 \times p\text{-}Akt^{T308} \times p\text{-}mTORC2^{S2481}$ |
| Dephosphorylation of the serine residue from p-$Akt^{T308, S473}$ | p-$Akt^{T308,S473}$ → p-$Akt^{T308}$ | $r32 = k32 \times p\text{-}Akt^{T308,S473}$ |
| Phosphorylation of Akt at the T308 residue to form the dual-phosphorylated species | p-$Akt^{S473}$ → p-$Akt^{T308,S473}$ | $r33 = k33 \times Akt^{S473} \times p\text{-}PDK1$ |
| Dephosphorylation of the threonine residue from p-$Akt^{T308, S473}$ | p-$Akt^{T308,S473}$ → p-$Akt^{S473}$ | $r34 = k34 \times p\text{-}Akt^{T308,S473}$ |
| Phosphorylation of the TSC complex via p-$Akt^{T308, S473}$ | $TSC_{clx}$ → p-$TSC_{clx}$ | $r35 = k35 \times TSC_{clx} \times p\text{-}Akt^{T308,S473}$ |
| Phosphorylation of the TSC complex via p-$Akt^{T308}$ | $TSC_{clx}$ → p-$TSC_{clx}$ | $r36 = k36 \times TSC_{clx} \times p\text{-}Akt^{T308}$ |
| Dephosphorylation of p-TSC complex | p-$TSC_{clx}$ → $TSC_{clx}$ | $r37 = k37 \times p\text{-}TSC_{clx}$ |
| Inactivation of mTORC1 via TSC | $mTORC1_{active}$ → $mTORC1_{inactive}$ | $r38 = k38 \times mTORC1_{active} \times TSC_{clx}$ |
| Activation of mTORC1 via leucine stimulation | $mTORC1_{inactive}$ → $mTORC1_{active}$ | $r39 = k39 \times mTORC1_{inactive} \times Leucine_{intracellular}$ |
| Phosphorylation of p70S6K1 via mTORC1 | p70S6K1 → p-p70S6K1$^{T389}$ | $r40 = k40 \times p70S6K1 \times mTORC1_{active}$ |

| | | |
|---|---|---|
| Phosphorylation of p70S6K1 via p-PDK1 | p70S6K1 → p-p70S6K1$^{T389}$ | $r41 = k41 \times p70S6K1 \times p\text{-}PDK1$ |
| Dephosphorylation of p-p70S6K1$^{T389}$ | p-p70S6K1$^{T389}$ → p70S6K1 | $r42 = k42 \times p\text{-}p70S6K1^{T389}$ |
| Inactivation of mTORC1 via p-p70S6K1$^{T389}$ | mTORC1$_{active}$ → mTORC1$_{inactive}$ | $r43 = k43 \times mTORC1_{active} \times p\text{-}p70S6K1^{T389}$ |
| Phosphorylation of PI3K variant | PI3K$_{variant}$ → p-PI3K$_{variant}$ | $r44 = k44 \times PI3K_{variant} \times p\text{-}IR\text{-}\beta^{Y1164}$ |
| Dephosphorylation of p-PI3K variant | p-PI3K$_{variant}$ → PI3K$_{variant}$ | $r45 = k45 \times p\text{-}PI3K_{variant}$ |
| Phosphorylation of mTORC2 | mTORC2 → p-mTORC2$^{S2481}$ | $r46 = k46 \times mTORC2 \times p\text{-}PI3K_{variant}$ |
| Dephosphorylation of mTORC2 | p-mTORC2$^{S2481}$ → mTORC2$^{S2481}$ | $r47 = k47 \times p\text{-}mTORC2^{S2481}$ |
| *Insulin oscillatory module (Sturis et al, 1991)** | | |
| Effect of glucose on insulin secretion | | $f_1(Glucose_{plasma}) = \dfrac{R_m}{1 + \exp\left(\dfrac{C_1 - \dfrac{Glucose_{plasma}}{V_g}}{a_1}\right)}$ |
| Effect of insulin on glucose production | | $f_2(Glucose_{plasma}) = U_b \cdot \left(1 - \exp\left(\dfrac{-Glucose_{plasma}}{C_2 \cdot V_g}\right)\right)$ |
| Insulin independent glucose utilization | | $f_3(Glucose_{plasma}) = \dfrac{Glucose_{plasma}}{C_3 \cdot V_g}$ |

| | | |
|---|---|---|
| Insulin dependent glucose utilization | | $f_4(Insulin_{interstitial}) = U_o + \frac{U_m - U_o}{1 + exp\left(-\beta \cdot ln\left(\frac{Insulin_{interstitial}}{C_4} \cdot \left(\frac{1}{V_i} + \frac{1}{E \cdot t_i}\right)\right)\right)}$ |
| Insulin dependent hepatic glucose production | | $f_5(x_3) = \frac{R_g}{1 + \exp\left(\alpha \cdot \left(\frac{x_3}{V_p - C_5}\right)\right)}$ |

*The insulin oscillatory module reaction equations are as presented for the 'Original Insulin-Glucose Feedback Model' in Tolic et al.[36]. These equations are simplified versions of the original equations presented in Sturis et al.[37], such that they were more straightforward to replicate in our model.

**Table S3. Related to Figure 2. System of ordinary differential equations.**

| Variable | Differential equations (with respect to time) |
|---|---|
| Leucine in the stomach | dx(1) = -r1 |
| Leucine in the gut | dx(2) = r1 – r2 – r3 – r68 |
| Plasma insulin | dx(3) = $f_1(Glucose_{plasma}) - E \cdot \left(\frac{Insulin_{plasma}}{V_p} - \frac{Insulin_{interstitial}}{V_i}\right) - \frac{Insulin_{plasma}}{t_p}$ |
| Plasma leucine | Equilibrium: dx(4) = r3 + r7 – r6 + $Leucine_{infusion}$<br>Simulation: dx(4) = r3 + r7 – r6 |
| Plasma KIC | dx(5) = r13 – r12 |
| Intracellular leucine | dx(6) = r6 + r9 + r11 – r7 – r10 – r15 |
| Intracellular KIC | dx(7) = r10 + r12 – r11 – r13 – r14 |
| Protein | dx(8) = r15 – r9 |
| IR-$\beta$ | dx(9) = r18 – r16 |
| p-IR-$\beta^{Y1164}$ | dx(10) = r16 – r17 |
| p-IR-$\beta_{refractory}$ | dx(11) = r17 – r18 |
| IRS1 | dx(12) = r20 + r22 – r19 – r21 |
| p-IRS1$^{Y}$ | dx(13) = r19 + r24 – r20 – r23 |
| p-IRS1$^{S}$ | dx(14) = r21 – r22 |
| PI3K | dx(15) = r24 – r23 |
| p-IRS1$^{Y}$_PI3K$_{clx}$ | dx(16) = r23 – r24 |
| PDK1 | dx(17) = r26 – r25 |
| p-PDK1 | dx(18) = r25 – r26 |
| Akt | dx(19) = r28 + r30 – r27 – r29 |
| p-Akt$^{T308}$ | dx(20) = r27 + r32 – r28 – r31 |
| p-Akt$^{S473}$ | dx(21) = r29 + r34 – r30 – r33 |

| | |
|---|---|
| p-Akt$^{T308,S473}$ | dx(22) = r31 + r33 – r32 – r34 |
| $TSC_{clx}$ | dx(23) = r37 – r35 – r36 |
| p-$TSC_{clx}$ | dx(24) = r35 + r36 – r37 |
| $mTORC1_{inactive}$ | dx(25) = r38 + r43 – r39 |
| $mTORC1_{active}$ | dx(26) = r39 – r38 – r43 |
| p70S6K1 | dx(27) = r42 – r40 – r41 |
| p-p70S6K1$^{T389}$ | dx(28) = r40 + r41 – r42 |
| $PI3K_{variant}$ | dx(29) = r45 – r44 |
| p-$PI3K_{variant}$ | dx(30) = r44 – r45 |
| mTORC2 | dx(31) = r47 – r46 |
| p-mTORC2$^{S2481}$ | dx(32) = r46 – r47 |
| Intercellular insulin | dx(35) = $E \cdot \left(\frac{Insulin_{plasma}}{V_p} - \frac{Insulin_{interstitial}}{V_i}\right) - \frac{Insulin_{interstitial}}{t_i}$ |
| Plasma glucose | dx(36) = $Glucose_{infusion} - f_2(Glucose_{plasma}) - f_3(Glucose_{plasma}) \cdot f_3(Insulin_{interstitial}) + f_5(x_3)$ |
| x1 | dx(37) = $\frac{3}{t_d} \cdot (Insulin_{insterstitial} - x_1)$ |
| x2 | dx(38) = $\frac{3}{t_d} \cdot (x_1 - x_2)$ |
| x3 | dx(39) = $\frac{3}{t_d} \cdot (x_2 - x_3)$ |

**Table S4. Related to Figure 2. Model species and initial concentrations.**
Model species (state variables) and their initial concentrations before and after the model calibration and at the end of the model equilibrium period.

| Speci-es | Name | Pre-calibration Value (M) | Calibrated Value (M) | Basis for values | | | Post model equilibrium (M) |
|---|---|---|---|---|---|---|---|
| | | | | Ref | Calc | Calib | |
| *Digestive module* | | | | | | | |
| $x(1)$ | Leucine in the stomach | 0 or 'leucine input at $t_x$'* | 0 | ☐ | ☐ | ☐ | 0 |
| $x(2)$ | Leucine in the gut | 0 | 0 | ☐ | ☐ | ☐ | 0 |
| *Leucine kinetic module* | | | | | | | |
| $x(4)$ | Plasma leucine | $1.21 \times 10^{-4}$ [1] | $1.21 \times 10^{-4}$ | ☒ | ☐ | ☐ | $1.21 \times 10^{-4}$ |
| $x(5)$ | Plasma KIC | $2.58 \times 10^{-5}$ [38] | $2.58 \times 10^{-5}$ | ☒ | ☐ | ☐ | $1.19 \times 10^{-5}$ |
| $x(6)$ | Intracellular leucine | $1.51 \times 10^{-4}$ [39] | $1.23 \times 10^{-4}$ | ☒ | ☐ | ☒ | $3.11 \times 10^{-6}$ |
| $x(7)$ | Intracellular KIC | N/a | $1.15 \times 10^{-5}$ | ☐ | ☐ | ☒ | $5.42 \times 10^{-6}$ |
| $x(8)$ | Protein | $1.34 \times 10^{-1}$ [40] | $1.34 \times 10^{-1}$ | ☒ | ☒ | ☐ | $1.34 \times 10^{-1}$ |
| *mTOR signalling module* | | | | | | | |
| $x(9)$ | IR-$\beta$ | $0.25 \times 10^{-9}$ [41] – $50 \times 10^{-9}$ [42] | $8.41 \times 10^{-10}$ | ☒ | ☒ | ☐ | $1.04 \times 10^{-9}$ |
| $x(10)$ | p-IR-$\beta^{Y1164}$ | N/a** | $1.68 \times 10^{-10}$ | ☐ | ☒ | ☐ | $1.12 \times 10^{-10}$ |
| $x(11)$ | p-IR-$\beta_{refractory}$ | N/a** | $1.68 \times 10^{-10}$ | ☐ | ☒ | ☐ | $2.63 \times 10^{-11}$ |
| $x(12)$ | IRS1 | $0.50 \times 10^{-9}$ [41] – $0.76 \times 10^{-9}$ [43] | $2.06 \times 10^{-10}$ | ☒ | ☒ | ☐ | $1.32 \times 10^{-10}$ |
| $x(13)$ | p-IRS1$^{Y}$ | N/a** | $4.12 \times 10^{-11}$ | ☐ | ☒ | ☐ | $1.80 \times 10^{-11}$ |
| $x(14)$ | p-IRS1$^{S}$ | N/a** | $4.12 \times 10^{-11}$ | ☐ | ☒ | ☐ | $1.21 \times 10^{-10}$ |
| $x(15)$ | PI3K | $<0.21 \times 10^{-9}$ [43] – $1.2 \times 10^{-9}$ [41] | $2.24 \times 10^{-10}$ | ☒ | ☐ | ☐ | $2.07 \times 10^{-10}$ |
| $x(16)$ | p-IRS1$^{Y}$/PI3K$_{clx}$ | N/a** | $4.48 \times 10^{-11}$ | ☐ | ☒ | ☐ | $6.18 \times 10^{-11}$ |
| $x(17)$ | PDK1 | $0.83 \times 10^{-9}$ [43] – $9.6 \times 10^{-9}$ [41] | $8.41 \times 10^{-10}$ | ☒ | ☐ | ☐ | $6.98 \times 10^{-10}$ |
| $x(18)$ | p-PDK1 | N/a** | $1.68 \times 10^{-10}$ | ☐ | ☒ | ☐ | $3.10 \times 10^{-10}$ |
| $x(19)$ | Akt | $0.39 \times 10^{-9}$ [43] – $49 \times 10^{-9}$ [41], $10 \times 10^{-9}$ [44] | $1.00 \times 10^{-8}$ | ☒ | ☐ | ☐ | $1.09 \times 10^{-8}$ |
| $x(20)$ | p-Akt$^{T308}$ | N/a** | $2.00 \times 10^{-9}$ | ☐ | ☒ | ☐ | $1.01 \times 10^{-9}$ |
| $x(21)$ | p-Akt$^{S473}$ | N/a** | $2.00 \times 10^{-9}$ | ☐ | ☒ | ☐ | $3.41 \times 10^{-9}$ |
| $x(22)$ | p-Akt$^{T308,S473}$ | N/a** | $2.00 \times 10^{-9}$ | ☐ | ☒ | ☐ | $7.07 \times 10^{-10}$ |

| | | | | | | | |
|---|---|---|---|---|---|---|---|
| $x(23)$ | $TSC_{clx}$ | $<0.21 \times 10^{-9}$ [43] – $5.2/11 \times 10^{-9}$ [41] | $6.94 \times 10^{-10}$ | ☒ | ☒ | ☐ | $5.79 \times 10^{-10}$ |
| $x(24)$ | p-$TSC_{clx}$ | N/a** | $1.39 \times 10^{-10}$ | ☐ | ☒ | ☐ | $2.53 \times 10^{-10}$ |
| $x(25)$ | $mTORC1_{inactive}$ | $4.3 \times 10^{-9}$ (mTOR)[41] / $5.4 \times 10^{-9}$ (RPTOR)[41] | $3.00 \times 10^{-10}$ | ☒ | ☒ | ☐ | $3.52 \times 10^{-10}$ |
| $x(26)$ | $mTORC1_{active}$ | N/a** | $6.00 \times 10^{-11}$ | ☐ | ☒ | ☐ | $8.02 \times 10^{-12}$ |
| $x(27)$ | p70S6K1 | $1.9 \times 10^{-9}$ [43] – $7.7 \times 10^{-9}$ [41] | $8.82 \times 10^{-9}$ | ☒ | ☒ | ☐ | $8.46 \times 10^{-9}$ |
| $x(28)$ | p-$p70S6K1^{T389}$ | N/a** | $1.76 \times 10^{-9}$ | ☐ | ☒ | ☐ | $2.12 \times 10^{-9}$ |
| $x(29)$ | $PI3K_{variant}$ | N/a | $2.06 \times 10^{-10}$ | ☐ | ☒ | ☐ | $2.35 \times 10^{-10}$ |
| $x(30)$ | p-$PI3K_{variant}$ | N/a | $4.11 \times 10^{-11}$ | ☐ | ☒ | ☐ | $1.18 \times 10^{-11}$ |
| $x(31)$ | mTORC2 | $4.3 \times 10^{-9}$ (RICTOR)[41] | $4.31 \times 10^{-10}$ | ☒ | ☒ | ☐ | $4.57 \times 10^{-10}$ |
| $x(32)$ | p-$mTORC2^{S2481}$ | N/a** | $8.63 \times 10^{-11}$ | ☐ | ☒ | ☐ | $6.03 \times 10^{-11}$ |
| *Insulin oscillatory module*[37] | | | | | | | |
| $x(3)$ | Plasma insulin | $2.80 \times 10^{-11}$ [45] | $2.80 \times 10^{-11}$ | ☒ | ☐ | ☐ | $2.71 \times 10^{-11}$ |
| $x(35)$ | Intercellular insulin | N/a*** | 0 | ☐ | ☐ | ☐ | $7.87 \times 10^{-12}$ |
| $x(36)$ | Plasma glucose | N/a*** | 0 | ☐ | ☐ | ☐ | $2.4 \times 10^{-3}$ |
| $x(37)$ | x1 | N/a*** | 0 | ☐ | ☐ | ☐ | $4.93 \times 10^{-12}$ |
| $x(38)$ | x2 | N/a*** | 0 | ☐ | ☐ | ☐ | $4.85 \times 10^{-12}$ |
| $x(39)$ | x3 | N/a*** | 0 | ☐ | ☐ | ☐ | $4.77 \times 10^{-12}$ |

Calc, calculations; Calib, calibrations; Ref, references

*Leucine input at $t_x$ corresponds to the leucine dose and timing that corresponds to the calibration and validation simulation.

**We were unable to locate concentrations of phosphorylated proteins, such that we assumed the initial concentrations of the phospho-proteins were 20% of the respective non-phosphorylated protein.

***Sturis et al.[37] did not provide initial concentrations for the respective species. We set the initial value to zero and allowed the model to achieve a steady-state value during the model equilibrium period.

**Table S5. Related to Figure 2. Model parameter values.**
Model parameter values before and after model calibration.

| Parameter name | Original | | Pre-calib. Value | Calibrated Value | Units | Basis for value | | | |
|---|---|---|---|---|---|---|---|---|---|
| | Value | Units | | | | Ref | Calc | Calib | Est^ |
| *Digestive module* | | | | | | | | | |
| $k_1$ | N/a* | | | $5.91\times10^{-1}$ | $min^{-1}$ | ☒ | ☐ | ☐ | ☒[46–48] |
| $k_2$ | N/a* | | | $1.86\times10^{-3}$ | $min^{-1}$ | ☒ | ☐ | ☐ | ☒[46–48] |
| $k_{68}$ | N/a* | | | $4.98\times10^{-3}$ | $min^{-1}$ | ☒ | ☐ | ☐ | ☒[46–48] |
| $k_3$ | N/a* | | | $1.11\times10^{-2}$ | $min^{-1}$ | ☒ | ☐ | ☐ | ☒[46–48] |
| *Blood Plasma* | | | | | | | | | |
| $k_4$ | N/a[#] | | $1.80\times10^{-8}$ | $2.01\times10^{-8}$ | $min^{-1}$ | ☐ | ☐ | ☒ | ☒ |
| *Leucine kinetic module* | | | | | | | | | |
| $k_6$ | $1.61\times10^{-7}$ [49] | $mol\cdot min^{-1}\cdot 100mL^{-1}$ | $1.73\times10^{-3}$ | $3.14\times10^{-2}$ | $min^{-1}$ | ☒ | ☒ | ☒ | ☐ |
| $k_7$ | $1.60\times10^{-7}$ [49] | $mol\cdot min^{-1}\cdot 100mL^{-1}$ | $1.73\times10^{-3}$ | $6.01\times10^{-4}$ | $min^{-1}$ | ☒ | ☒ | ☒ | ☐ |
| $k_{15}$ | $1.30\times10^{-7}$ [49] | $mol\cdot min^{-1}\cdot 100mL^{-1}$ | $1.40\times10^{-3}$ | $4.44\times10^{4}$ | $mol^{-1}\cdot min^{-1}$ | ☒ | ☒ | ☒ | ☐ |
| $k_9$ | $1.49\times10^{-7}$ [49] | $mol\cdot min^{-1}\cdot 100mL^{-1}$ | $4.54\times10^{-3}$ | $2.23\times10^{-15}$ | $mol\cdot min^{-1}$ | ☒ | ☒ | ☒ | ☐ |
| $k_{10}$ | $9.53\times10^{-8}$ [49] | $mol\cdot min^{-1}\cdot 100mL^{-1}$ | $1.03\times10^{-3}$ | $1.18\times10^{-2}$ | $min^{-1}$ | ☒ | ☒ | ☒ | ☐ |
| $k_{11}$ | $7.35\times10^{-8}$ [49] | $mol\cdot min^{-1}\cdot 100mL^{-1}$ | $5.17\times10^{-3}$ | $2.93\times10^{-2}$ | $min^{-1}$ | ☒ | ☒ | ☒ | ☐ |
| $k_{12}$ | N/a[#] | | $5.19\times10^{-3}$ | $5.33\times10^{-2}$ | $min^{-1}$ | ☒ | ☐ | ☒ | ☒[50] |
| $k_{13}$ | N/a[#] | | $9.69\times10^{-4}$ | $1.90\times10^{-2}$ | $min^{-1}$ | ☒ | ☐ | ☒ | ☒[50] |
| $k_{14}$ | $1.96\times10^{-8}$ [49] | $mol\cdot min^{-1}\cdot 100mL^{-1}$ | $1.38\times10^{-3}$ | $9.56\times10^{-2}$ | $min^{-1}$ | ☒ | ☒ | ☒ | ☐ |
| *mTOR signalling module* | | | | | | | | | |
| $k_{16}$ | 0.0254 [51] | A.U. | $3.32\times10^{8}$ | $3.37\times10^{7}$ | $mol^{-1}\cdot min^{-1}$ | ☒ | ☐ | ☒ | ☒ |
| $k_{17}$ | 0.149 [51] | A.U. | $1.56\times10^{-1}$ | $7.27\times10^{-2}$ | $min^{-1}$ | ☒ | ☐ | ☒ | ☒ |
| $k_{18}$ | 0.0310 [51] | A.U. | $6.87\times10^{-3}$ | $2.93\times10^{-1}$ | $min^{-1}$ | ☒ | ☐ | ☒ | ☒ |

| | | | | | | | | | |
|---|---|---|---|---|---|---|---|---|---|
| $k_{19}$ | 0.135 [51] | A.U. | $2.66\times10^{7}$ | $9.57\times10^{6}$ | mol$^{-1}$· min$^{-1}$ | ☒ | ☐ | ☒ | ☒ |
| $k_{20}$ | $3.28\times10^{-3}$ [51] | A.U. | $1.04\times10^{-1}$ | $2.14\times10^{-1}$ | min$^{-1}$ | ☒ | ☐ | ☒ | ☒ |
| $k_{21}$ | 1 [51] | A.U. | $2.04\times10^{5}$ | $5.78\times10^{4}$ | mol$^{-1}$· min$^{-1}$ | ☒ | ☐ | ☒ | ☒ |
| $k_{22}$ | $1.00\times10^{-4}$ [51] | A.U. | $1.20\times10^{-2}$ | $6.88\times10^{-3}$ | min$^{-1}$ | ☒ | ☐ | ☒ | ☒ |
| $k_{23}$ | $7.06\times10^{12}$ [52] | min$^{-1}$ | $3.19\times10^{8}$ | $1.65\times10^{8}$ | mol$^{-1}$· min$^{-1}$ | ☒ | ☒ | ☒ | ☒ |
| $k_{24}$ | 10 [52] | min$^{-1}$ | $1.58\times10^{-1}$ | $1.97\times10^{-1}$ | min$^{-1}$ | ☒ | ☒ | ☒ | ☒ |
| $k_{25}$ | N/a[#] | | $4.22\times10^{8}$ | $2.93\times10^{8}$ | mol$^{-1}$· min$^{-1}$ | ☐ | ☐ | ☒ | ☒ |
| $k_{26}$ | N/a[#] | | 1.92 | $4.05\times10^{-1}$ | min$^{-1}$ | ☐ | ☐ | ☒ | ☒ |
| $k_{27}$ | 0.700 [51] | A.U. | $5.74\times10^{8}$ | $2.26\times10^{8}$ | mol$^{-1}$· min$^{-1}$ | ☒ | ☐ | ☒ | ☒ |
| $k_{28}$ | 4.07 [51] | A.U. | $4.45\times10^{1}$ | $5.03\times10^{1}$ | min$^{-1}$ | ☒ | ☐ | ☒ | ☒ |
| $k_{29}$ | N/a[#] | | $5.15\times10^{8}$ | $2.73\times10^{9}$ | mol$^{-1}$· min$^{-1}$ | ☐ | ☐ | ☒ | ☒ |
| $k_{30}$ | N/a[#] | | $1.69\times10^{1}$ | $1.03\times10^{1}$ | min$^{-1}$ | ☐ | ☐ | ☒ | ☒ |
| $k_{31}$ | 4.51 [51] | A.U. | $3.33\times10^{12}$ | $2.90\times10^{12}$ | mol$^{-1}$· min$^{-1}$ | ☒ | ☐ | ☒ | ☒ |
| $k_{32}$ | 7.53 [51] | A.U. | $7.06\times10^{3}$ | $6.28\times10^{3}$ | min$^{-1}$ | ☒ | ☐ | ☒ | ☒ |
| $k_{33}$ | N/a[#] | | $6.53\times10^{7}$ | $2.30\times10^{7}$ | mol$^{-1}$· min$^{-1}$ | ☐ | ☐ | ☒ | ☒ |
| $k_{34}$ | N/a[#] | | $5.00\times10^{1}$ | $6.47\times10^{-1}$ | min$^{-1}$ | ☐ | ☐ | ☒ | ☒ |
| $k_{35}$ | $1.00\times10^{-4}$ [51] | A.U. | $4.99\times10^{4}$ | $2.13\times10^{4}$ | mol$^{-1}$· min$^{-1}$ | ☒ | ☐ | ☒ | ☒ |
| $k_{36}$ | $6.27\times10^{-3}$ [51] | A.U. | $7.73\times10^{5}$ | $9.31\times10^{5}$ | mol$^{-1}$· min$^{-1}$ | ☒ | ☐ | ☒ | ☒ |
| $k_{37}$ | $8.13\times10^{-3}$ [51] | A.U. | $1.06\times10^{-1}$ | $8.47\times10^{-2}$ | min$^{-1}$ | ☒ | ☐ | ☒ | ☒ |
| $k_{38}$ | 1.00 [51] | A.U. | $5.18\times10^{5}$ | $2.04\times10^{5}$ | mol$^{-1}$· min$^{-1}$ | ☒ | ☐ | ☒ | ☒ |
| $k_{39}$ | 0.0514 [51] | A.U. | 2.64 | $3.36\times10^{-1}$ | mol$^{-1}$· min$^{-1}$ | ☒ | ☐ | ☒ | ☒ |
| $k_{40}$ | $5.74\times10^{-3}$ [51] | A.U. | $8.51\times10^{7}$ | $4.44\times10^{7}$ | mol$^{-1}$· min$^{-1}$ | ☒ | ☐ | ☒ | ☒ |
| $k_{41}$ | N/a[#] | | $3.61\times10^{6}$ | $8.14\times10^{4}$ | mol$^{-1}$· min$^{-1}$ | ☐ | ☐ | ☒ | ☒ |
| $k_{42}$ | $5.28\times10^{-3}$ [51] | A.U. | $3.93\times10^{-1}$ | $7.18\times10^{-2}$ | min$^{-1}$ | ☒ | ☐ | ☒ | ☒ |
| $k_{43}$ | N/a[#] | | $8.83\times10^{5}$ | $7.59\times10^{5}$ | mol$^{-1}$· min$^{-1}$ | ☐ | ☐ | ☒ | ☒ |

| | | | | | | | | | |
|---|---|---|---|---|---|---|---|---|---|
| $k_{44}$ | 1.00 [51] | A.U. | $9.34\times10^{6}$ | $1.64\times10^{6}$ | $mol^{-1}\cdot min^{-1}$ | ☒ | ☐ | ☒ | ☒ |
| $k_{45}$ | $2.32\times10^{-4}$ [51] | A.U. | $1.28\times10^{-1}$ | $1.02\times10^{-1}$ | $min^{-1}$ | ☒ | ☐ | ☒ | ☒ |
| $k_{46}$ | $3.19\times10^{-2}$ [51] | A.U. | $1.75\times10^{7}$ | $2.23\times10^{7}$ | $mol^{-1}\cdot min^{-1}$ | ☒ | ☐ | ☒ | ☒ |
| $k_{47}$ | $2.56\times10^{-2}$ [51] | A.U. | $6.97\times10^{-2}$ | $2.90\times10^{-2}$ | $min^{-1}$ | ☒ | ☐ | ☒ | ☒ |
| *Insulin oscillatory module* [37] | | | | | | | | | |
| $k_{48}$ ($V_p$) | 3 | L | 3 | 3 | L | ☒ | ☐ | ☐ | ☐ |
| $k_{49}$ ($V_i$) | 11 | L | 11 | 11 | L | ☒ | ☐ | ☐ | ☐ |
| $k_{50}$ ($V_g$) | 10 | L | 10 | 10 | L | ☒ | ☐ | ☐ | ☐ |
| $k_{51}$ (E) | 0.2 | $L\cdot min^{-1}$ | 0.2 | $9.55\times10^{-2}$ | $L\cdot min^{-1}$ | ☒ | ☐ | ☒ | ☐ |
| $k_{52}$ ($t_p$) | 6 | min | 6 | 3.62 | min | ☒ | ☐ | ☒ | ☐ |
| $k_{53}$ ($t_i$) | 100 | min | 100 | 92.6 | min | ☒ | ☐ | ☒ | ☐ |
| $k_{54}$ ($t_d$) | 36 | min | 36 | 28 | min | ☒ | ☐ | ☒ | ☐ |
| $k_{55}$ ($R_m$) | 210 | $mU\cdot min^{-1}$ | $1.40\times10^{-9}$ | $1.12\times10^{-9}$ | $mol\cdot min^{-1}$ | ☒ | ☒ | ☒ | ☐ |
| $k_{56}$ ($a_1$) | 300 | $mg\cdot L^{-1}$ | $1.67\times10^{-3}$ | $9.21\times10^{-4}$ | $mol\cdot L^{-1}$ | ☒ | ☒ | ☒ | ☐ |
| $k_{57}$ ($C_1$) | 2000 | $mg\cdot L^{-1}$ | $1.11\times10^{-2}$ | $9.61\times10^{-3}$ | $mol\cdot L^{-1}$ | ☒ | ☒ | ☒ | ☐ |
| $k_{58}$ ($U_b$) | 72 | $mg\cdot min^{-1}$ | $4.00\times10^{-4}$ | $2.70\times10^{-4}$ | $mol\cdot min^{-1}$ | ☒ | ☒ | ☒ | ☐ |
| $k_{59}$ ($C_2$) | 144 | $mg\cdot L^{-1}$ | $7.99\times10^{-4}$ | $6.04\times10^{-4}$ | $mol\cdot L^{-1}$ | ☒ | ☒ | ☒ | ☐ |
| $k_{60}$ ($C_3$) | 1000 | $mg\cdot L^{-1}$ | $5.55\times10^{-3}$ | $2.62\times10^{-3}$ | $mol\cdot L^{-1}$ | ☒ | ☒ | ☒ | ☐ |
| $k_{61}$ ($U_0$) | 40 | $mg\cdot min^{-1}$ | $2.22\times10^{-4}$ | $1.71\times10^{-4}$ | $mol\cdot min^{-1}$ | ☒ | ☒ | ☒ | ☐ |
| $k_{62}$ ($U_m$) | 940 | $mg\cdot min^{-1}$ | $5.22\times10^{-3}$ | $4.61\times10^{-3}$ | $mol\cdot min^{-1}$ | ☒ | ☒ | ☒ | ☐ |
| $k_{63}$ ($\beta$) | 1.77 | unitless | 1.77 | 1.33 | unitless | ☒ | ☐ | ☒ | ☐ |
| $k_{64}$ ($C_4$) | 80 | $mU\cdot L^{-1}$ | $5.34\times10^{-10}$ | $3.89\times10^{-10}$ | $mol\cdot L^{-1}$ | ☒ | ☒ | ☒ | ☐ |
| $k_{65}$ ($R_g$) | 180 | $mg\cdot min^{-1}$ | $9.99\times10^{-4}$ | $7.94\times10^{-4}$ | $mol\cdot min^{-1}$ | ☒ | ☒ | ☒ | ☐ |
| $k_{66}$ ($\alpha$) | 0.29 | $L\cdot mU^{-1}$ | $4.35\times10^{10}$ | $4.50\times10^{10}$ | $L\cdot mol^{-1}$ | ☒ | ☒ | ☒ | ☐ |
| $k_{67}$ ($C_5$) | 26 | $mU\cdot L^{-1}$ | $1.73\times10^{-10}$ | $1.39\times10^{-10}$ | $mol\cdot L^{-1}$ | ☒ | ☒ | ☒ | ☐ |

A.U., arbitrary units; Calc, calculations; Calib, calibrations; Est, estimations; N/a, not applicable; Ref, references

^Citations specific to the estimation column were used for estimation of the respective parameter.

*We manually derived the parameter values for the digestive system module using the plasma leucine time-course[1] and two published values. 1) True ileal digestibility[48]: approximately 90% of the ingested leucine is absorbed into the blood stream (i.e., 10% of the ingested leucine bolus is excreted). 2) First-pass splanchnic extraction[46,47]: in young adults, the first-pass splanchnic extraction of amino acids is 23-29%.

#Parameter values that have an original value of 'N/a' were added during the model topology expansion. These parameters did not have references to directly infer the parameter value, such that we used a heuristic approach to derive the pre-calibration values. Our heuristic approach required each parameter value to produce a reasonable qualitative kinetic for the respective model species. A reasonable qualitative kinetic for the model species was defined as: 1) does not remain at zero throughout the simulation, 2) is not fully saturated throughout the model simulation, and 3) has a reasonable dynamic following model simulation (i.e., there was a realistic increase and decrease in the species that corroborated with upstream species).

**Table S6. Related to Figure 2A. Calibration dataset characteristics.**

| Species | Study | Subjects (M, F) | Age (SEM) | Intervention | Leucine content | EAA content | Whey protein |
|---|---|---|---|---|---|---|---|
| Plasma leucine | Glynn et al.[1] | 7 (3, 4) | 29 (2) | A single bolus provided (noncaloric, noncaffeinated carbonated beverage, 500 mL) | 3.5 g | 10.0 g | 0 g |
| $F_{m,a}$ | | | | | | | |
| $F_{m,0}$ | | | | | | | |
| FSR | | | | | | | |
| Plasma insulin | | | | | | | |
| Intracellular leucine | Drummond et al.[4] | 7 (3, 4) | 29.0 (2.0) | A single bolus provided (in a 500 mL solution) | 3.5 g | 10.0 g | 0 g |
| p-Akt$^{S473}$ | Meta-analyzed | | | | | | |
| p-p70S6K$^{T389}$ | Meta-analyzed | | | | | | |

**Table S7. Related to Figure 2B,C. Validation dataset characteristics.**

| Study | Subjects (M, F) | Age (SEM) | Intervention | Leucine content | EAA content | Whey protein |
|---|---|---|---|---|---|---|
| Glynn et al.[1] | 7 (3, 4) | 32 (2) | Single bolus provided (noncaloric, noncaffeinated carbonated beverage, 500 mL) | 1.85 g | 10.0 g | 0 g |
| Mitchell et al.[2] | 8 (8, 0) | 19.7 (0.5) | Single bolus provided (in aqueous solution, 250 mL) | 3.59 g | 15.0 g | 0 g |
| Mitchell et al.[2] | 8 (8, 0) | 21.5 (1.1) | Four equal fractions were ingested at 45-min intervals (in aqueous solution, 250 mL) | 3.59 g (4 x 0.9 g) | 15.0 g | 0 g |
| Drummond et al.[4] | 7 (3, 4) | 29.0 (2.0) | Single bolus provided (in a 500 mL solution) | 3.5 g | 10.0 g | 0 g |
| Wilkinson et al.[3] | 7 (7,0) | 21 (0.3) | Leucine intervention: Single bolus provided (with ~400 mL of water) | 3.42 g | 3.42 g | 0 g |
| Dickinson et al.[5] | 8 (3,5) | 25 (2) | A 10-gram EAA solution was ingested. | 1.8 g | 10 g | 0 g |

**Table S8. Related to Figure 2. Amino acid profile for each intervention.**

| Dataset | Leu (g) | His (g) | Iso (g) | Lys (g) | Met (g) | Phe (g) | Thr (g) | Trp (g) | Val (g) | Total AA (g) |
|---|---|---|---|---|---|---|---|---|---|---|
| Glynn et al.[1] - leucine | 3.50 | 0.80 | 0.80 | 1.20 | 0.30 | 1.40 | 1.00 | 0 | 1.00 | 10.00 |
| Glynn et al.[1] - control | 1.85 | 1.10 | 1.00 | 1.55 | 0.30 | 1.55 | 1.45 | 0 | 1.20 | 10.00 |
| Mitchell et al.[2] - bolus | 3.59 | 1.21 | 1.73 | 3.07 | 0.95 | 0.91 | 0.48 | 1.13 | 1.86 | 14.93 |
| Mitchell et al.[2] - pulsatile | 3.59 | 1.21 | 1.73 | 3.07 | 0.95 | 0.91 | 0.48 | 1.13 | 1.86 | 14.93 |
| Drummond et al.[4] | 3.50 | 0.80 | 0.80 | 1.20 | 0.30 | 1.40 | 1.00 | 0 | 1.00 | 10.00 |
| Dickinson et al.[5] | 1.80 | 1.10 | 1.00 | 1.60 | 0.30 | 1.60 | 1.40 | 0 | 1.20 | 10.00 |
| Wilkinson et al.[3] | 3.42 | 0 | 0 | 0 | 0 | 0 | 0 | 0 | 0 | 3.42 |

*Atherton et al.[50], Moore et al.[7], and Mazzulla et al.[8] are not included in this table as each study provided participants with whey protein and did not report the amino acid profile of the intervention.

10. Bertuzzi, A., Conte, F., Mingrone, G., Papa, F., Salinari, S., and Sinisgalli, C. (2016). Insulin Signaling in Insulin Resistance States and Cancer: A Modeling Analysis. PloS one *11*, e0154415. 10.1371/journal.pone.0154415.

40. Wolfe, R.R., Park, S., Kim, I.-Y., Starck, C., Marquis, B.J., Ferrando, A.A., and Moughan, P.J. (2019). Quantifying the contribution of dietary protein to whole body protein kinetics: examination of the intrinsically labeled proteins method. American journal of physiology. Endocrinology and metabolism *317*, E74–E84. 10.1152/ajpendo.00294.2018.

41. Nagaraj, N., Wisniewski, J.R., Geiger, T., Cox, J., Kircher, M., Kelso, J., Pääbo, S., and Mann, M. (2011). Deep proteome and transcriptome mapping of a human cancer cell line. Molecular systems biology *7*, 548. 10.1038/msb.2011.81.

42. Varusai, T.M., and Nguyen, L.K. (2018). Dynamic modelling of the mTOR signalling network reveals complex emergent behaviours conferred by DEPTOR. Scientific reports *8*, 643. 10.1038/s41598-017-18400-z.

43. Beck, M., Schmidt, A., Malmstroem, J., Claassen, M., Ori, A., Szymborska, A., Herzog, F., Rinner, O., Ellenberg, J., and Aebersold, R. (2011). The quantitative proteome of a human cell line. Molecular systems biology *7*, 549. 10.1038/msb.2011.82.

44. Hatakeyama, M., Kimura, S., Naka, T., Kawasaki, T., Yumoto, N., Ichikawa, M., Kim, J.-H., Saito, K., Saeki, M., Shirouzu, M., et al. (2003). A computational model on the modulation of mitogen-activated protein kinase (MAPK) and Akt pathways in heregulin-induced ErbB signalling. Biochemical Journal *373*, 451–463. 10.1042/bj20021824.

45. Reidy, P.T., Walker, D.K., Dickinson, J.M., Gundermann, D.M., Drummond, M.J., Timmerman, K.L., Fry, C.S., Borack, M.S., Cope, M.B., Mukherjea, R., et al. (2013). Protein blend ingestion following resistance exercise promotes human muscle protein synthesis. The Journal of nutrition *143*, 410–416. 10.3945/jn.112.168021.

46. Boirie, Y., Gachon, P., and Beaufrère, B. (1997). Splanchnic and whole-body leucine kinetics in young and elderly men. The American journal of clinical nutrition *65*, 489–495. 10.1093/ajcn/65.2.489.

47. Volpi, E., Mittendorfer, B., Wolf, S.E., and Wolfe, R.R. (1999). Oral amino acids stimulate muscle protein anabolism in the elderly despite higher first-pass splanchnic extraction. The American journal of physiology *277*, E513-20. 10.1152/ajpendo.1999.277.3.E513.

48. Wolfe, R.R., Park, S., Kim, I.-Y., Starck, C., Marquis, B.J., Ferrando, A.A., and Moughan, P.J. (2019). Quantifying the contribution of dietary protein to whole body protein kinetics: examination of the intrinsically labeled proteins method. American journal of physiology. Endocrinology and metabolism *317*, E74–E84. 10.1152/ajpendo.00294.2018.

49. Tessari, P., Inchiostro, S., Zanetti, M., and Barazzoni, R. (1995). A model of skeletal muscle leucine kinetics measured across the human forearm. The American journal of physiology *269*, E127-36.

50. Atherton, P.J., Etheridge, T., Watt, P.W., Wilkinson, D., Selby, A., Rankin, D., Smith, K., and Rennie, M.J. (2010). Muscle full effect after oral protein: time-dependent concordance and discordance between human muscle protein synthesis and mTORC1 signaling. The American journal of clinical nutrition *92*, 1080–1088. 10.3945/ajcn.2010.29819.

51. Dalle Pezze, P., Sonntag, A.G., Thien, A., Prentzell, M.T., Gödel, M., Fischer, S., Neumann-Haefelin, E., Huber, T.B., Baumeister, R., Shanley, D.P., et al. (2012). A dynamic network model of mTOR signaling reveals TSC-independent mTORC2 regulation. Science Signaling *5*. 10.1126/scisignal.2002469.

52. Chew, Y.H., Shia, Y.L., Lee, C.T., Majid, F.A.A., Chua, L.S., Sarmidi, M.R., and Aziz, R.A. (2009). Modeling of glucose regulation and insulin-signaling pathways. Molecular and Cellular Endocrinology *303*, 13–24. 10.1016/j.mce.2009.01.018.
